# A multi-omics approach identifies pancreatic cancer cell extracellular vesicles as mediators of the unfolded protein response in normal pancreatic epithelial cells

**DOI:** 10.1101/2021.10.12.464079

**Authors:** Charles P. Hinzman, Shivani Bansal, Yaoxiang Li, Anton Iliuk, Michael Girgis, Baldev Singh, Kelly M. Herremans, Jose G. Trevino, Vijay K. Singh, Partha P. Banerjee, Amrita K. Cheema

## Abstract

Although cancer-derived extracellular vesicles (cEVs) are thought to play a pivotal role in promoting cancer progression events, their precise effect on neighboring normal cells is unknown. In this study, we investigated the impact of pancreatic cancer ductal adenocarcinoma (PDAC) derived EVs on recipient non-tumorigenic pancreatic normal epithelial cells upon internalization. We show that PDAC cEVs increase the proliferation and invasive capability of treated normal cells. We further demonstrate that cEVs induce endoplasmic reticulum (ER) stress and the unfolded protein response (UPR) in treated normal pancreatic epithelial cells within 24 hours. Subsequently, these cells release several inflammatory cytokines. Leveraging a layered multi-omics approach, we analyzed EV cargo from a panel of 6 PDAC and 2 normal pancreas cell lines, using multiple EV isolation methods. We found that cEVs were enriched for an array of biomolecules which can induce or regulate ER stress and the UPR, including palmitic acid, sphingomyelins, metabolic regulators of tRNA charging and proteins which regulate trafficking and degradation. We further show that palmitic acid, at doses relevant to those found in cEVs, is sufficient to induce ER stress in normal pancreas cells. These results suggest that cEV cargo packaging may be designed to disseminate proliferative and invasive characteristics upon internalization by distant recipient normal cells, hitherto unreported. This study is among the first to highlight a major role for PDAC cEVs to induce stress in treated normal pancreas cells that may modulate a systemic response leading to altered phenotypes. For the first time, our study implicates cEV transported palmitic acid as a potential driver in this process. These findings highlight the importance of EVs in mediating disease etiology and open potential areas of investigation toward understanding the role of cEV lipids in promoting cell transformation in the surrounding microenvironment.

## Introduction

Pancreatic cancer is projected to become the 2^nd^ leading cause of cancer-related mortality in the United States and Europe by 2030^1,2^. The estimated 5-year survival rate currently remains ∼9%. Two key drivers of these poor outcomes are increasing incidence (the number of patients diagnosed each year has doubled over the last 20 years) and late-stage diagnosis^3^. Pancreatic ductal adenocarcinoma (PDAC) is the most common type of pancreatic cancer. Unfortunately, 80% of patients with this lethal malignancy are ineligible for surgical resection due to late-stage disease at diagnosis^4,5^. Though key genetic alterations are known to coincide with disease progression, this knowledge has not led to the development of early detection assays^6^. A key feature in PDAC is the diversity of cell types in the tumor microenvironment; as much as 90% of the tumor volume can consist of non-cancer cells^7^. In recent years, several studies have sought to understand the role of various cell types in the tumor microenvironment and the interaction between PDAC tumor cells and other cell populations, including cancer associated fibroblasts (CAFs)^8-10^, immune cells^11-13^ and endothelial cells^14,15^. Importantly, complex intracellular communication is needed to facilitate coordinated efforts that are conducive to tumor growth. While these studies demonstrate how different cell types may harmonize with cancer cells for disease progression, many of these are focused on very late stages of disease, often when patients have few treatment options. A better understanding of mediators of early formation of this tumor microenvironment is critical for improving clinical outcomes.

Extracellular vesicles (EVs) are nanometer sized lipid-bilayer particles released from cells of all tissue types^16,17^. EVs play important signal mediating roles in a variety of normal physiological processes, as well as in several pathologies, including cancer^18-21^. EV cargo is a rich source of potential biomarkers for PDAC diagnosis^22-24^ and can mediate survival signaling between CAFs and cancer cells^25,26^. EVs also suppress the immune system in the tumor microenvironment and can prime metastatic sites for pre-cancer cell infiltration^27-29^. Although these studies suggest that EVs likely regulate intracellular communication within the late-stage tumor microenvironment, few studies have attempted to address the role of EVs in mediating earliest stages of PDAC development. Herein, we tested the hypothesis that PDAC EVs could mediate distinct biochemical, genetic, or metabolic alterations in treated normal pancreatic epithelial cells. Using a systematic multi-pronged omics approach, we show that PDAC cell-derived EVs (cEVs) induce a myriad of stress response pathways in treated normal pancreatic epithelial cells. These events culminate in activation of the unfolded protein response, driving inflammation and altered cell behavior. These studies highlight a novel stress activation paradigm by which cEVs may prime the microenvironment driving disease onset and tumor progression.

## Results

### Key differences exist between extracellular vesicles isolated from pancreas cancer cell lines compared to normal pancreas cell lines

In order to validate our EV isolation methods, we characterized our sample preparations using orthogonal techniques including immunoblot analysis, cryogenic electron microscopy (Cryo-EM) and nanoparticle tracking analysis (NTA). One of the striking observations was the differential expression between certain “canonical” EV markers between pancreas cancer and normal cells. EVs isolated from the normal cell line hTERT-HPNE, for example, did not express the protein Programmed cell death 6-interacting protein (PDCD6IP), also known as ALG-2-interacting protein (ALIX) (**Fig. 1a**). Cargo sorting into small EVs involves many proteins within the endosomal sorting complex required for transport (ESCRT) pathway, such as ALIX, and thus it is commonly used as a marker for exosomes^30^. Furthermore, EVs isolated from the cancer cell line MiaPaCa-2 did not express epithelial cell adhesion molecule (EPCAM) (**Suppl. Fig. S1**), another widely used marker for exosomes. We also observed, via NTA, that pancreatic cancer cells secreted on average 2-8-fold more EVs than normal pancreatic cells (**Fig. 1b**), a finding we confirmed by cryo-EM (**Suppl. Fig. S2**). Finally, NTA showed that EVs isolated from all the 8 cell lines used in our study differed in terms of average particle diameter, ranging from ∼144 nm in HPDE-H6c7 cells to ∼325 nm in Capan-1 cells. (**Suppl. Fig. S1**), an initial observation we continue to investigate. These results highlight the importance of characterizing EV samples isolated from various cell culture lines using multiple approaches, to identify key differences in EV populations.

**Figure 1:**
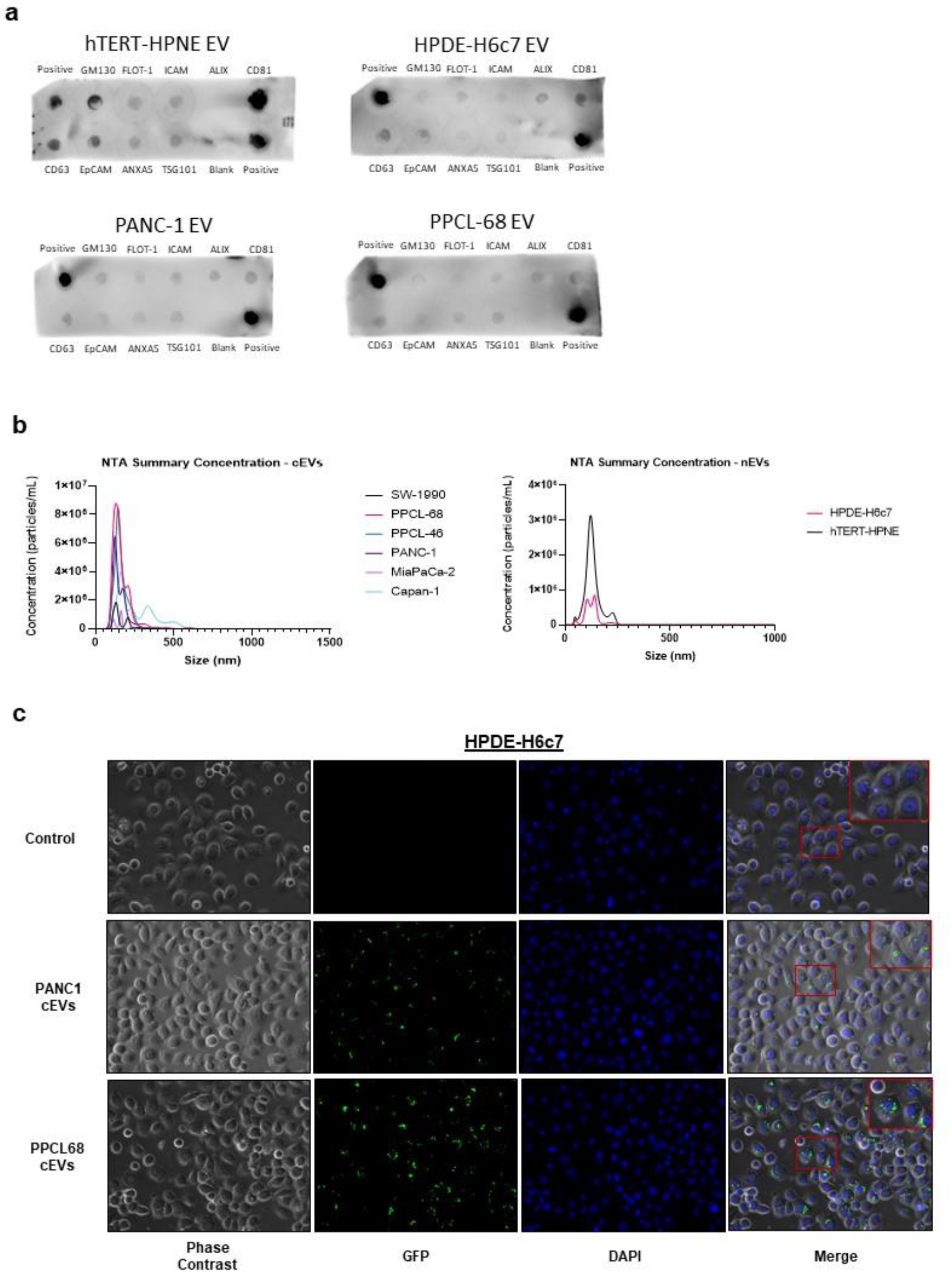
Extracellular vesicles derived from pancreas cancer cells significantly differ from those isolated from normal pancreas cells. **a**. Immunoblot arrays comparing the expression of known EV markers in EVs isolated from the normal pancreas cell lines hTERT-HPNE, HPDE-H6c7 and the pancreas cancer cell lines PANC-1 and PPCL-68. **b**. Nanoparticle tracking analysis (NTA) concentration data of cancer-cell EVs (cEVs, left) and normal-cell EVs (nEVs, right) demonstrating key differences in EV yield. **c**. Representative phase contrast and fluorescent microscopy images of the normal pancreas cell line HPDE-H6c7 incubated for 18-24 hours with cEVs from either PANC-1 or PPCL-68 cells. cEVs were labeled with green fluorescence using PKH67 dye prior to administration to HPDE-H6c7 cells.

### Pancreatic cancer cell EVs induce significant gene expression changes in treated normal pancreatic epithelial cells

We asked if pancreatic cancer-cell-derived EVs (cEVs) mediate altered signaling in cEV treated normal pancreatic epithelial cells. We first performed EV internalization experiments, to ensure that cEVs were internalized by cEV treated normal cells. We found that both cEVs isolated from PANC-1 and PPCL-68 cells were readily absorbed by HPDE-H6c7 and hTERT-HPNE cells (**Fig. 1c, Suppl. Fig. S1**). To identify potential changes, we used RNA-Seq to investigate global gene expression changes in normal cells post-cEV treatment. Using cEVs isolated from PANC-1 and PPCL-68 (a patient-derived xenograft (PDX) cell line^31^) PDAC cells, we treated two normal cell line models, hTERT-HPNE and HPDE-H6c7 (*n* = 3 per group), with cEVs or a negative control (residual media left behind after enrichment of cEVs by ultracentrifugation) for 24 hours (**Fig. 2a**). Visualization of gene expression using MA plots indicated robust change in the global gene expression profiles of normal cells post cEV treatment (**Fig. 2b**).

**Figure 2:**
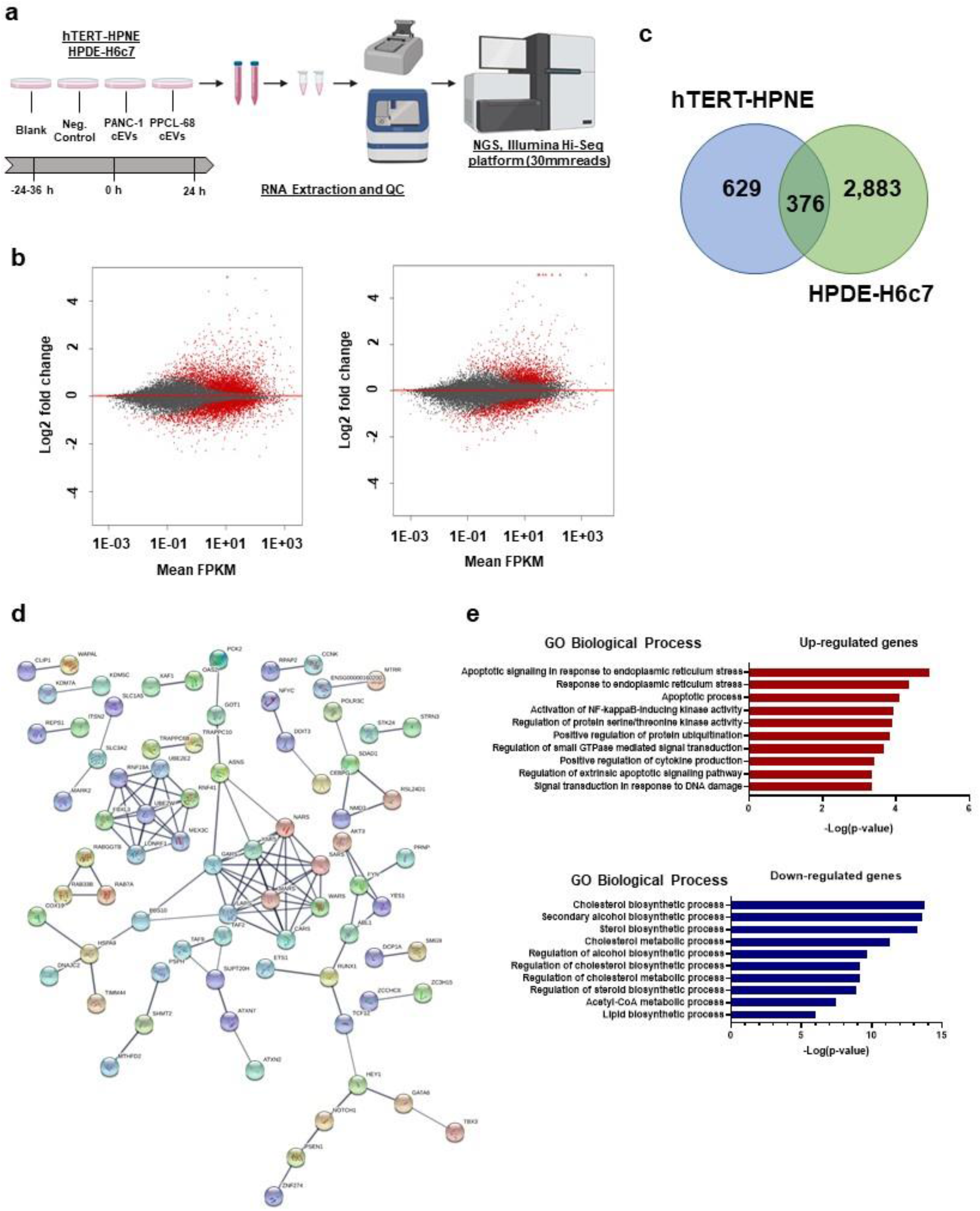
Pancreas cancer cell extracellular vesicles (cEVs) induce robust gene expression changes in treated normal pancreas cells. **a**. Experimental design for RNA-Seq experiment on hTERT-HPNE and HPDE-H6c7 cells treated with PANC-1 or PPCL-68 derived cEVs. **b**. MA plots showing robust gene expression changes in HPDE-H6c7 (top) and hTERT-HPNE (bottom) cells post-24-hour treatment with PANC-1 cEVs. Red dots indicate genes which are significantly dysregulated (FDR-adj. *p* < 0.05 and fold-change <0.5 or >2, *n* = 3 per condition). **c**. Number of significantly differentially expressed genes in hTERT-HPNE and HPDE-H6c7 cells after 24-hour treatment with cEVs. **d**. STRING analysis visualizing protein-protein interactions between upregulated common DEGs in hTERT-HPNE and HPDE-H6c7 cells (PPI-enrichment *p-*value = 2.09e-07). Line thickness represents confidence of the interaction and only high-confidence interactions are shown. Unconnected nodes are hidden. This network was enriched for genes in the aminoacyl-tRNA biosynthesis (adj. *p* = 2.69e-05), cytosolic tRNA aminoacylation (adj. *p* = 1.19e-06) and ATF4 (adj. *p* = 0.0402)/ATF6 (adj. *p* = 0.0402) pathways. **e**. Gene ontology (GO) biological process classification of the 376 common DEGs between hTERT-HPNE and HPDE-H6c7 cells. These genes were enriched in biological processes regulating endoplasmic reticulum (ER) stress, NF-kB signaling, cytokine production and response to DNA damage, while lipid synthesis was severely altered, with decreased expression in genes regulating several cholesterol, sterol and acetyl-CoA synthesis pathways were severely altered.

Gene set enrichment analysis (GSEA) identified 1,005 and 3,259 differentially expressed genes (DEGs) (FDR-adj. *p* < 0.05, see methods) in hTERT-HPNE and HPDE-H6c7 cells, respectively (**Fig. 2c, Suppl. Data 1**). We found 376 DEGs commonly dysregulated in both cell lines in response to both cEV treatments, but not in the negative control group. Next, we performed protein-protein interaction network analysis using the Search Tool for the Retrieval of Interacting Genes/Proteins (STRING) database^32,33^. The 252 upregulated DEGs within this gene set revealed significant protein-protein interactions (PPI enrichment *p =* 2.09e-07) (**Fig. 2d**). This network contained genes in the aminoacyl-tRNA biosynthesis (adj. *p* = 2.69e-05), cytosolic tRNA aminoacylation (adj. *p* = 1.19e-06) and ATF4 (adj. *p* = 0.0402)/ATF6 (adj. *p* = 0.0402) pathways. Interestingly, the remaining downregulated DEGs similarly exhibited significantly increased protein-protein interaction (PPI enrichment *p* = 0.00528) particularly for genes involved in protein refolding (adj. *p* = 0.0087) (**Suppl. Fig S3**). Gene ontology (GO) analysis of the 376 common DEGs identified enrichment in biological processes associated with endoplasmic reticulum (ER) stress, nuclear factor kappa-light-chain-enhancer of activated B cells (NF-κB) signaling, cytokine production and response to DNA damage (**Fig. 2f, Suppl. Data 2**). Lipid synthesis was significantly dysregulated, with decreased expression in genes regulating cholesterol, sterol, and acetyl-CoA synthesis pathways (**Fig. 2f, Suppl. Data 2**). Taken together, these findings suggest that normal pancreatic cells undergo distinct stress responses after cEV treatment, altering the biosynthesis of lipid species.

Though widely used as normal pancreatic cell models, hTERT-HPNE and HPDE-H6c7 lines differ in origin, method of immortalization, and morphology. We analyzed DEGs uniquely dysregulated in each cell line in order to examine cell line specific responses to cEV treatment. In HPDE-H6c7 cells, a model better representing a true epithelial cell, pathway analysis revealed enrichment in the unfolded protein response (UPR) and ER stress signaling, inositol metabolism and adipocytokine signaling (**Suppl. Fig. S3, Suppl. Data 3**), whereas cell cycle pathways were down-regulated (**Suppl. Fig. S3, Suppl. Data 3**). Interestingly, in hTERT-HPNE cells, an intermediate acinar-to-ductal metaplasia state cell model, we again found that cEVs increased expression of genes regulating UPR/ER stress (including TNF Receptor Associated Factor 6 (*TRAF6*)-mediated Interferon Regulatory Factor 7 (*IRF7*) activation), but also hippo, tumor necrosis factor-alpha (TNF-α) and interferon-gamma (IFN-γ) signaling (**Suppl. Fig. S3, Suppl. Data 3)**. Lipid biosynthesis, transforming growth factor-beta (TGF-β)-dependent extracellular matrix regulation and NOTCH signaling were down-regulated (**Suppl. Fig. S3, Suppl. Data 3**). These results highlight that individual cell lines differentially respond to cEV treatment, though the induction of UPR/ER stress seemed to be a consistent theme between these models underscoring the biological relevance and significance of this pathway perturbation in the treated cells.

### cEVs induce the unfolded protein response and ER stress in treated normal pancreatic epithelial cells

Given that the UPR/ER stress pathway was predominant in the GO analysis of both common and unique DEGs, and implicated in our STRING analysis, this warranted further investigation. Motif analysis using the Hypergeometric Optimization of Motif EnRichment (HOMER) platform^34^ showed the DEG gene set preferentially contained target motifs of C/EBP homologous protein (*CHOP)*, also known as DNA damage-inducible transcript 3 (*DDIT3*) **Fig. 3a)**. Additional transcription factor analysis using the Enrichr platform^35,36^ confirmed this observation (**Fig. 3b**). CHOP is a key transcription factor upregulated in response to elevated UPR^37,38^. Transcriptional regulatory network analysis further identified overrepresentation of key regulatory elements targeted by several transcription factors responsible for regulating UPR/ER stress, including X-Box Binding Protein 1 (*XBP1), DDIT3*, Aryl Hydrocarbon Receptor Nuclear Translocator *(ARNT)*, Hypoxia Inducible Factor 1 Subunit Alpha *(HIF1A)*, and Activating Transcription Factors 6, 4 and 3 *(ATF6, ATF4 and ATF3)* (**Fig. 3b, Suppl. Fig. S4**). In the UPR/ER stress pathways, the leading genes enriched in our dataset (*DDIT3*, Heat Shock Protein Family A (Hsp70) Member 5 *(HSPA5)*, Nuclear Transcription Factor Y Subunit Beta *(NFYB)*, Nuclear Transcription Factor Y Subunit Gamma *(NFYC), ATF6* and *ATF4)* were upregulated post cEV treatment (**Fig. 3c**) suggesting cEVs induce the UPR in treated normal cells.

**Figure 3:**
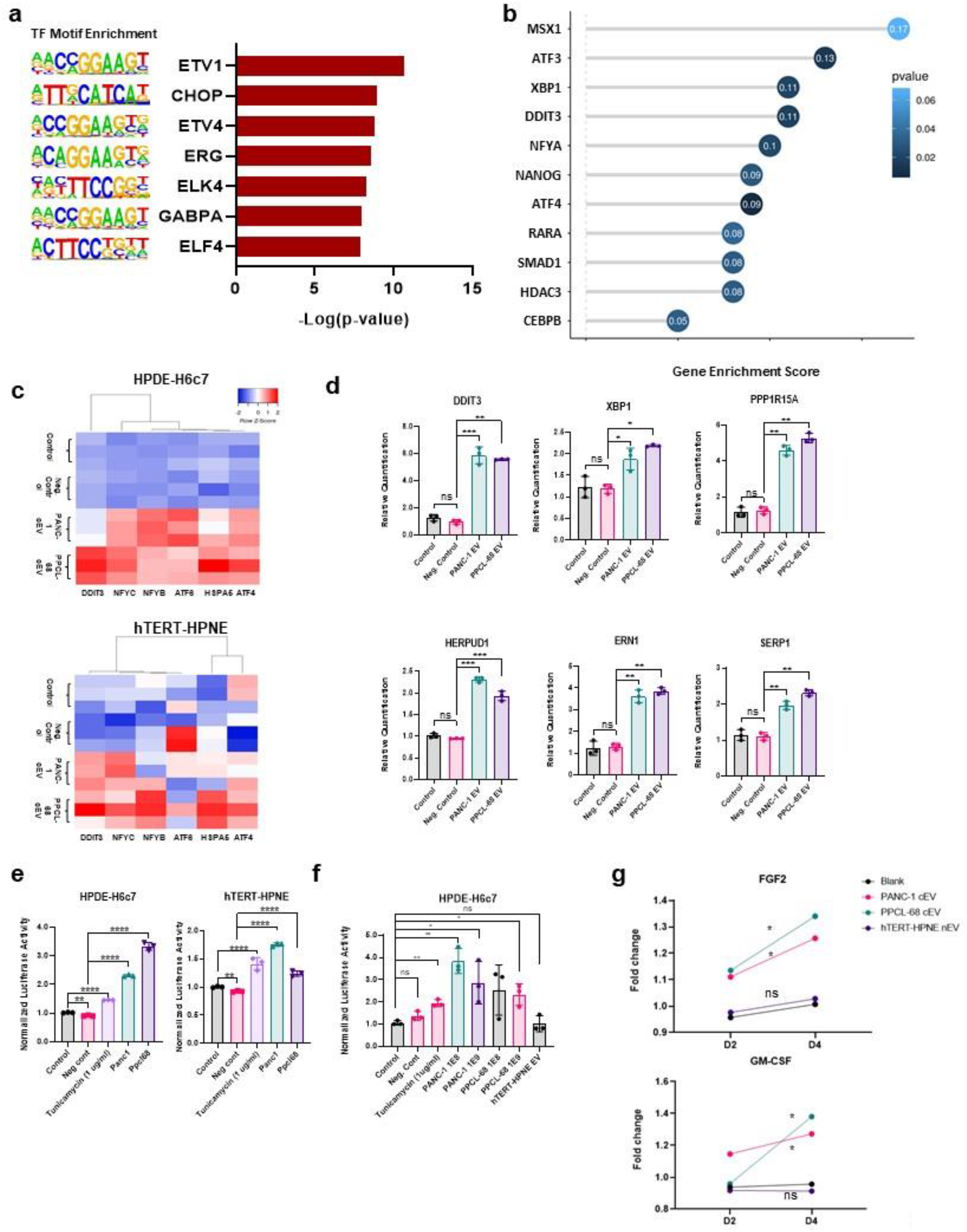
cEVs induce unfolded protein response/ER stress in treated normal pancreatic epithelial cells. **a**. Select known motifs identified using HOMER as significantly enriched (*p*-value < 0.05) in DEGs **b**. Transcription Regulatory Relationships Unraveled by Sentence-based Text Mining (TRRUST) transcriptional regulatory network analysis results, predicting transcription factors responsible for regulating genes enriched in our gene set. Select results shown from HPDE-H6c7 upregulated genes. **c**. Heatmap showing leading genes in the UPR/ER Stress pathways, enriched in our DEG gene set in HPDE-H6c7 (top) and hTERT-HPNE (bottom). *N* = 3 per condition. **d**. qPCR data recapitulating upregulation in key mediators of UPR/ER stress in HPDE-H6c7 cells treated with cEVs for 24 hours. *N* = 3 per condition. **e**. Luciferase quantification showing upregulation in binding activity of the ER stress-response element (ERSE) promoter, indicating increased UPR/ER stress in cEV treated normal cells 24 hours post treatment. Tunicamycin serves as a positive control for UPR/ER Stress induction. *N* = 3 per condition. **f**. Luciferase quantification data showing upregulation in binding activity of ERSE promoter, induced by treatment with cEVs isolated using size exclusion chromatography. *N* = 3 per condition. **g**. Cytokine quantification data showing regulation of fibroblast growth factor 2 (FGF2) and granulocyte-macrophage colony-stimulating factor (GM-CSF) in the media of cEV treated normal cells treated with cEVs after 2 and 4 days. *N* = 4 per condition. ns = *p* > 0.05, * = *p* ≤ 0.05, ** = *p* ≤ 0.01, *** = *p* ≤ 0.001 and **** = *p* ≤ 0.0001.

Next, we validated these findings using qPCR. In independent experiments, we treated both normal cell lines (hTERT-HPNE and HPDE-H6c7) with separate batches of cEVs derived from PANC-1 or PPCL-68 PDAC cells, as well as negative control (residual media from the cEV isolation process). We confirmed up-regulation of key UPR and ER stress regulators including *XBP1, DDIT3*, Endoplasmic Reticulum to Nucleus Signaling 1 *(ERN1)*, Protein phosphatase 1 regulatory subunit 15A (*PPP1R15A)*, also known as Growth Arrest and DNA-Damage-Inducible 34 (*GADD34)*, Homocysteine Inducible ER Protein with Ubiquitin Like Domain 1 *(HERPUD1)* and Stress Associated Endoplasmic Reticulum Protein 1 *(SERP1)* (**Fig. 3d**). We further validated ER stress induction using hTERT-HPNE and HPDE-H6c7 cell lines stably transfected with an ER Stress Response Element (ERSE) luciferase reporter. After cEV treatment, there was significant upregulation in ERSE-reporter activity in both cell lines (**Fig. 3e**). Importantly, this effect was recapitulated when treating cells with cEVs isolated using a separate size-exclusion chromatography (SEC)-based method, but not by treatment with EVs isolated from normal hTERT-HPNE cells (**Fig. 3f**). We further observed ER stress induction using a lower dose of cEVs, persisting 48 hours post treatment (**Fig. 3f, Suppl. Fig S5**). This suggests that UPR/ER stress induction is independent of cEV isolation method and is specific to pancreatic cancer-derived cell EVs and that this response is EV dose dependent.

Since prolonged ER stress and the UPR can induce inflammation, we performed a time course experiment to see if cEV treated normal cells released inflammatory cytokines. We found that after 2 and 4 days post-cEV treatment, treated normal cells secreted significantly higher levels of fibroblast growth factor 2 (FGF2) and granulocyte-macrophage colony-stimulating factor (GM-CSF) (**Fig. 3g**). However, treatment with nEVs isolated from the normal cell line, hTERT-HPNE, did not result in increased cytokine release, suggesting this response is specific to cEVs (**Fig. 3g**). Prolonged ER stress can also induce apoptosis; therefore, we next investigated whether treated normal cells underwent increased apoptosis. We examined cell death by measuring caspase 3/7 activity, live/dead cell staining and observing nuclei number and morphology using DAPI staining. We did not observe signs of apoptosis in cEV treated cells, indicating that UPR/ER stress in this context is not leading to cell death (**Suppl. Fig S5**). Since UPR/ER stress has been shown to impact cell proliferation^39^, we asked if cEVs influenced cell number. Hence, we treated normal cells with cEVs and measured cell number by both DAPI and crystal violet staining. Strikingly, we found that cEV treatment of normal pancreatic epithelial cells significantly increased the cell number suggesting that the ER stress induced by cEV treatment resulted in increased cell proliferation. (**Suppl. Fig. S6**).

Taken together, these findings confirm that cEV treatment can cause ER stress and activate the UPR in normal cells, validating our initial observation with RNA-Seq analysis. Furthermore, cEVs increased the proliferative capability of normal cells, did not induce apoptosis, and finally caused an inflammatory response in treated normal cells. Taken together, these data demonstrate that cEV-mediated ER stress may be reprogramming normal cell function, potentially, a key step in aiding alterations in cellular phenotype that may lead to microenvironmental remodeling conducive to disease progression.

### Proteomics, lipidomics and metabolomics studies reveal enrichment of potential UPR/ER stress mediators in cEVs

The novel finding that cEV treated normal cells show remarkable changes in gene expression and increased proliferative capability has striking implications on molecular changes that may eventually lead to early onset of neoplastic transformation. To better understand this phenomenon, we sought to identify potential mediators of these changes contained in the EV cargo. We performed multi-omics analysis and quantification of the biomolecular content contained in cEVs derived from 6 different pancreatic cancer cell lines (PANC-1, SW-1990, Capan-1, MiaPaCa-2, PPCL-68, and PPCL-46) and compared their biochemical profiles to normal cell EVs (nEVs) derived from the normal cell lines hTERT-HPNE and HPDE-H6c7 (**Fig. 4a**). We used two independent EV isolation methods, ultracentrifugation with filtration (UC) and EVTrap, a magnetic bead-based isolation method, for proteomics profiling experiments to obviate potential protein contamination issues introduced by a given EV enrichment method. Principal Component Analysis (PCA) of proteomics and targeted metabolomics profiling revealed clear separation between cEVs and nEVs, indicating distinct protein, small molecule, and lipid profiles (**Fig. 4b-e**). Interestingly, each approach identified an array of distinct molecules enriched in cEV cargo that could potentially mediate and/or regulate UPR/ER stress in normal epithelial cells.

**Figure 4:**
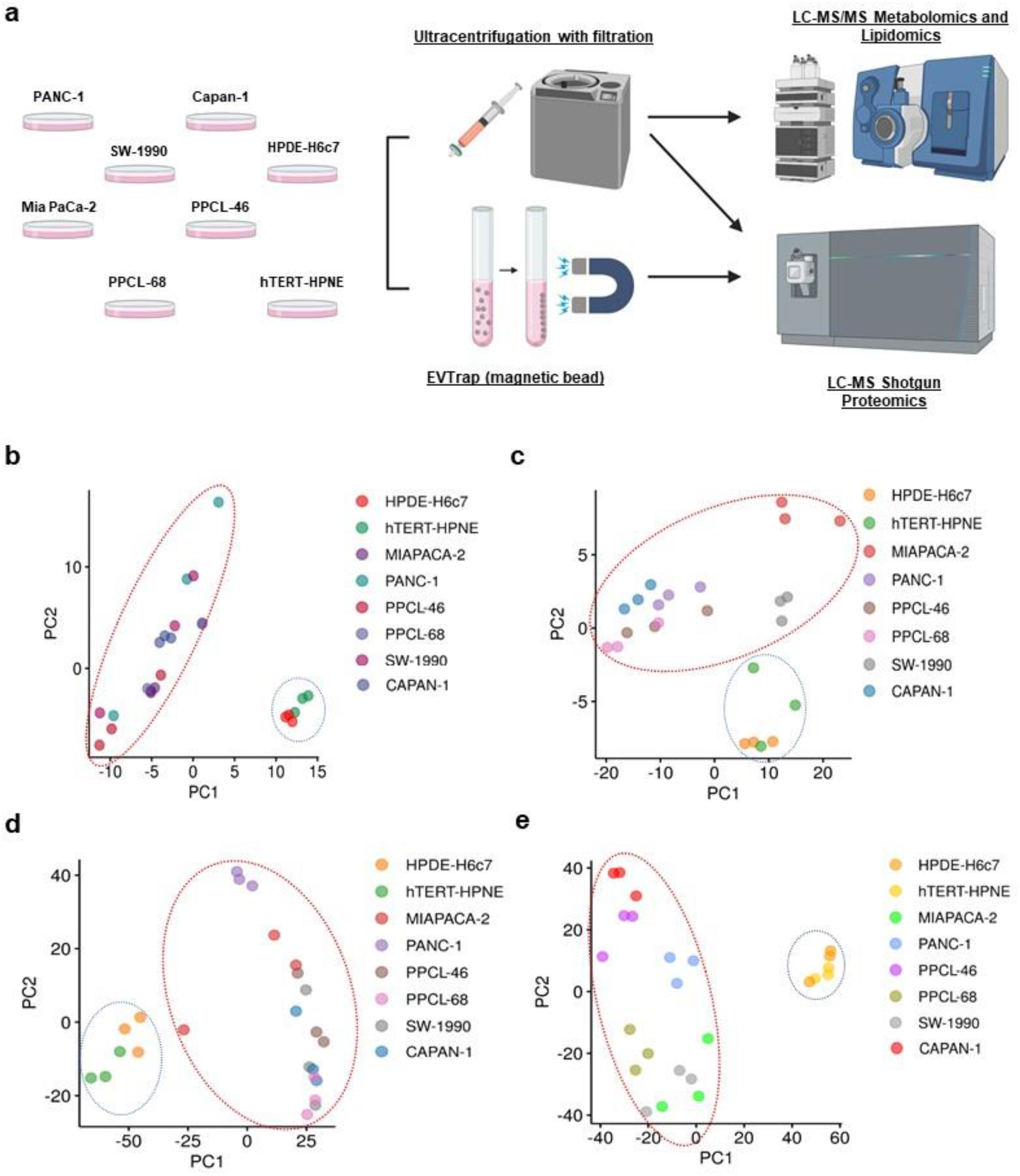
Multi-omics analysis of cEVs and nEVs reveals enrichment in potential regulators of UPR/ER stress. **a**. Experimental protocol for EV isolation from 6 pancreatic cancer cell lines (cEV) and 2 normal pancreas cell line (nEV) models. Proteomics was performed on EVs isolated using two independent methods, ultracentrifugation and EVTrap. **b-e**. Principal component analysis (PCA) plots of **b**. polar metabolites **c**. lipids and **d**. proteins isolated by EVTrap and **e**. proteins isolated by ultracentrifugation, in cEVs and nEVs. Red circles encompass cEV samples, blue circles encompass nEV samples. *N* = 3 independent EV isolations per cell line.

Shotgun proteomics revealed 332 proteins significantly dysregulated in cEVs compared to nEVs between both ultracentrifugation and EVTrap isolation methods (**Fig. 5a**). Of these, several of the upregulated proteins are involved in targeted protein degradation and ubiquitination, including tumor susceptibility 101 (TSG101), member RAS oncogene family (RAB40C), UBA domain containing 2 (UBAC2), cullin 5 (CUL5), RAS like proto-oncogene A (RALA), and transmembrane protein 59 (TMEM59) (**Fig. 5b**). We validated upregulation of TSG101, TMEM59, UBAC2, RALA and RAB40C in cEVs using immunoblot (**Fig. 5c**). We posit that cEVs enrichment for these specific proteins could likely alter the normal protein degradation process, impacting protein folding and clearance functions, when internalized by normal cells.

**Figure 5:**
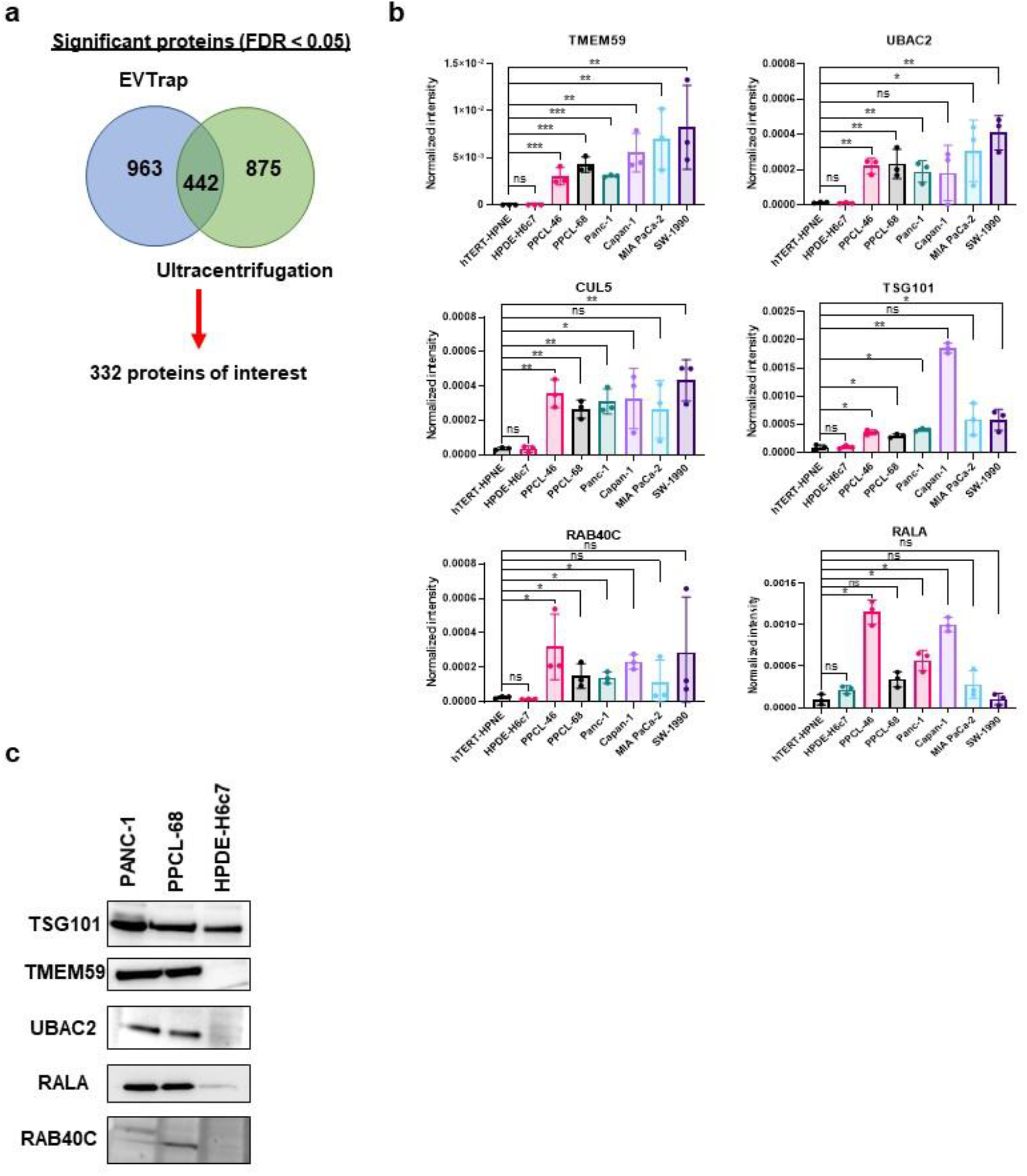
Shotgun proteomics reveals potential mediators of UPR/ER stress are enriched in cEVs. **a**. Venn diagram showing number of significantly dysregulated proteins identified by either EVTrap or ultracentrifugation EV isolation methods. The total number of proteins yielded 332 proteins of interest, defined as FDR-adjusted *p*-value < 0.05, Fold change > 3 or < 0.5, and a matching expression trend (either upregulated or downregulated) in each isolation method (EVTrap and ultracentrifugation). **b**. Select proteins involved in the regulation of protein transport and ubiquitination, as defined by gene ontology analysis and literature search, which were enriched in cEVs. Values are plotted as means with standard deviation. Representative intensity data from EV samples isolated by ultracentrifugation. *N* = 3 independent EV isolations per cell line. **c**. Immunoblot validations of TSG101, TMEM59, UBAC2, RALA and RAB40C from proteomics data. Images captured from independent EV isolations from each cell line. ns = *p* > 0.05, * = *p* ≤ 0.05, ** = *p* ≤ 0.01, and *** = *p* ≤ 0.001.

Separately, we also performed multiple reaction monitoring (MRM)-based quantitative mass spectrometry, which included analysis of 269 polar metabolites and 1,021 lipid species separately, that provided novel insights into potential mediators of UPR/ER stress enriched in cEV cargo. Visualization of significantly dysregulated lipids using a heatmap, confirmed differential expression patterns between cEVs and nEVs (**Fig. 6a**). Further investigation identified enrichment of lipid species containing a palmitic acid (16:0) moiety, as well as several sphingomyelin (SM) species in cEVs (**Fig. 6b**). Palmitic acid has been shown to induce UPR in pancreatic islet cells ^40,41^ while altered sphingolipid metabolism is a contributor to the UPR^42,43^. To investigate whether this could be occurring in our cEV treated normal cells, we treated HPDE-H6c7 ERSE luciferase reporter cells with various concentrations of palmitic acid. We found that ER stress was induced at 75 μM (**Fig. 6c**), well below commonly reported doses of 250 μM - 500 μM and higher in other cell types^44,45^. This suggests that enrichment of palmitic acid in cEVs may induce ER stress in part, leading to activation of the UPR in treated normal cells.

**Figure 6:**
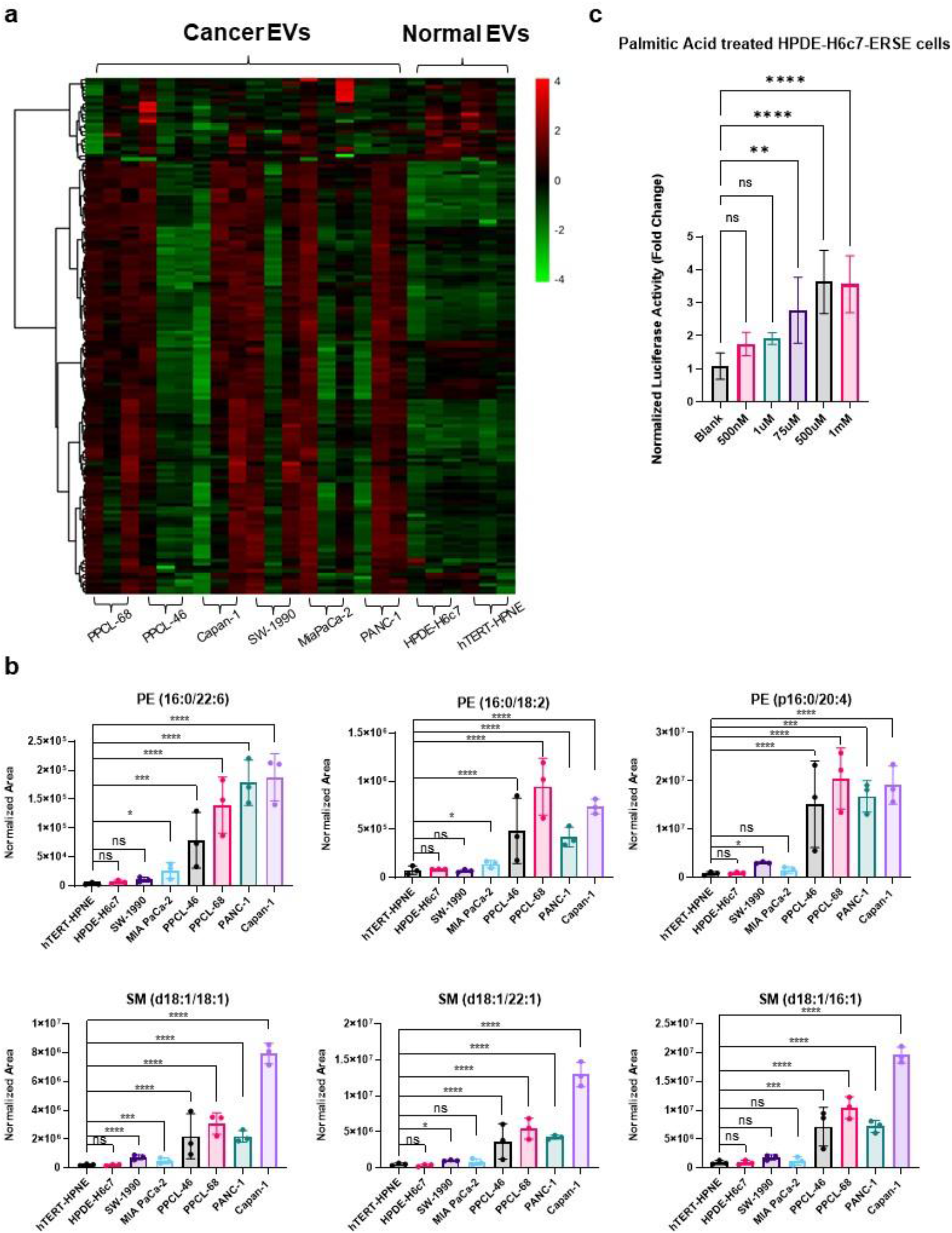
cEVs and nEVs have distinct lipid profiles. **a**. Heatmap depicting distinct lipid expression profiles in cEVs compared to nEVs, as determined by LC-MS/MS. **b**. Select lipid species containing palmitic acid (16:0) and SM lipid species which were significantly upregulated in cEVs. Means with standard deviation are plotted. *N* = 3 independent EV isolations per cell line. **c**. ERSE-luciferase activity plotted from HPDE-H6c7 cells treated with various concentrations of palmitic acid esterified to bovine serum albumin. Means with standard deviation are plotted. *N* = 5 per condition. ns = *p* > 0.05, * = *p* ≤ 0.05, ** = *p* ≤ 0.01, and *** = *p* ≤ 0.001.

Ingenuity Pathway Analysis (IPA) of significantly dysregulated metabolites in cEV cargo showed over representation of the citrulline biosynthesis and tRNA charging pathways (**Fig. 7a**). This aligned with our previous STRING and RNA-Seq analysis of unique DEGs. Metabolites central to these processes, including arginine, ornithine, N-acetylornithine, glutamine, and proline, were significantly dysregulated in cEVs compared to nEVs (**Suppl. Data 4**). Interestingly, the amino acids arginine, glutamine, and proline were all significantly downregulated in cEVs compared to nEVs (**Fig. 7b**). N-acetylornithine was also downregulated, whereas cEVs were enriched for ornithine (**Suppl. Data 4**). We also observed enrichment of oncometabolites, such as succinate, and accumulation of phenylalanine in cEVs (**Suppl. Data 4, Fig. 7b**). We validated upregulation of ornithine, succinate, phenylalanine, and downregulation of arginine in separate MRM-mass spectrometry-based validation experiments (**Suppl. Fig. S7**). Taken together, our data, for the first time demonstrate that cEVs are enriched for multiple classes of biomolecules which may synergistically induce UPR/ER in normal pancreatic cells that may drive some of the earliest molecular signaling changes in normal pancreatic cells.

**Figure 7:**
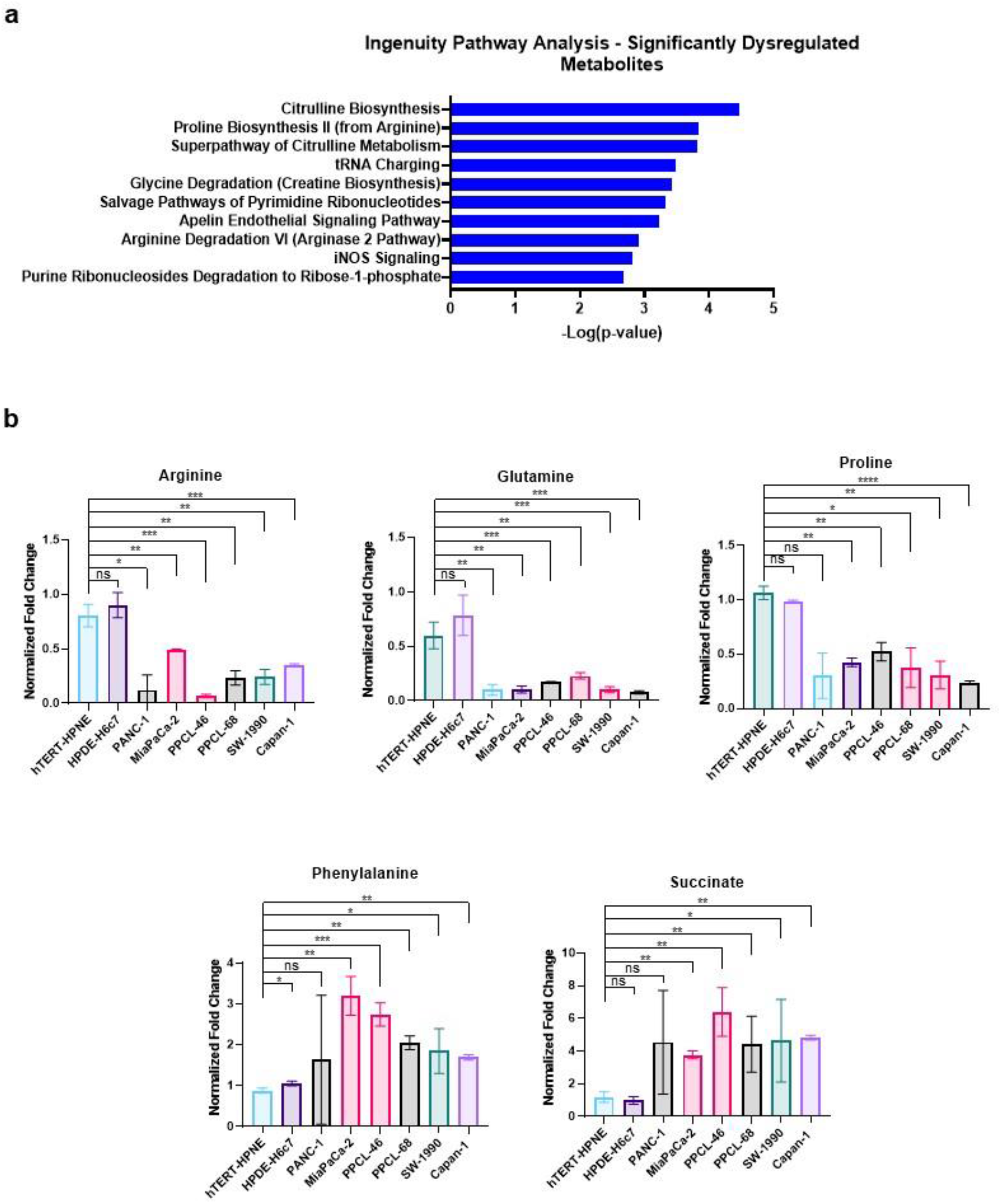
cEVs are enriched for oncometabolites and molecules which alter tRNA aminoacylation in treated normal pancreas cells. **a**. Ingenuity pathway analysis (IPA) identified significant dysregulation in metabolites related to citrulline biosynthetic pathways and tRNA charging in cEVs compared to nEVs. **b**. Levels of arginine, glutamine, proline, phenylalanine, and succinate in cEVs and nEVs as determined by LC-MS/MS. Means with standard deviations are plotted. *N* = 3 independent EV isolations per condition. ns = *p* > 0.05, * = *p* ≤ 0.05, ** = *p* ≤ 0.01, *** = *p* ≤ 0.001 and **** = *p* ≤ 0.0001.

## Discussion

While pancreatic cancer cell derived EVs (cEVs) are thought to regulate intercellular communication and signal transduction leading to proliferation, invasion and metastasis, a deeper understanding of how these events are mediated is critical to understand disease etiology. How specific molecular signaling events, triggered upon cEV internalization by normal cells, impact the function of those normal cells remains understudied. Especially in the context of pancreatic cancer, understanding key changes within non-cancerous cells that shape the microenvironment is critical toward designing novel therapeutic interventions and diagnostics. In this study, using an array of human PDAC and normal pancreas cell line models, we have demonstrated that pancreatic cancer cEVs drastically alter the behavior and gene expression profiles of treated normal pancreatic cells. These changes are dramatic, and our findings elucidate a new perspective for understanding pancreas cancer onset and progression.

Specifically, our data show that cEVs cause ER stress and activate the unfolded protein response (UPR) in treated cells. Subsequently, this leads to an inflammatory response and drives increased proliferation. Furthermore, we demonstrate that cEVs are enriched for an array of molecular mediators of ER stress and the UPR, including protein homeostasis regulators, oncometabolites and palmitic acid. These changes are dramatic in the context of expected normal cell behavior, and thus warrant further investigation as mediators of disease progression. The endoplasmic reticulum is crucial for protein maturation and proper folding, with approximately 30% of all proteins undergoing ER processing^46,47^. If this intricate process is disrupted, inducing ER stress, cells respond with coping mechanisms that include the UPR^46^. Canonically under sustained ER-stress, apoptosis is triggered leading to cell clearance^48^. Importantly, our data demonstrate that apoptosis is not triggered in these cells up to 48 hours post-cEV treatment, and in fact, cEVs significantly increased the proliferative capability of treated cells. This suggests a possible survival mechanism is activated in these cells.

Given the fundamental role of the UPR in normal physiology, abnormal ER stress has also been implicated across an array of pathologies including cancer. IRE1α is the fifth-most mutated kinase in cancer and the UPR has been shown to impact many cancer hallmarks, including angiogenesis, genome stability, inflammation, metastasis, and drug resistance^49,50^. Specific to the pancreas, UPR is constitutively active due to high demand for protein synthesis and maintenance of acinar cell homeostasis^51^. However, prolonged UPR can lead to inflammation and as such, UPR is a driver in chronic pancreatitis and diabetes^52,53^. Our data demonstrate that cEVs induce an inflammatory response, with upregulation of both FGF2 and GM-CSF. ER stress further mediates the progression of pancreatic intraepithelial neoplasia (PanIN) to pancreatic ductal adenocarcinoma (PDAC) through expression of anterior gradient-2 (AGR2)^54^.

Framing our findings within the context of these previous reports, it is possible that in the earliest stages of PDAC development, cancer cells release EVs which are internalized by neighboring normal cells, inducing ER stress, promoting inflammation and disease progression.

As we continue to investigate potential mediators of these signaling changes, it seems clear that analyzing multiple classes of EV biomolecules is important. Each of our analyses identified potential inducers of ER stress. Our proteomics analysis of EVs indicated significant enrichment of proteins which are integrally involved with protein trafficking and degradation. An interesting finding was that TSG101 is enriched in pancreatic cEVs; TSG101 is widely used as a marker for demonstrating enrichment of EVs generally, given its association with the endosomal sorting complex required for transport (ESCRT) pathway^30^. Though TSG101 functions in the biogenesis of multivesicular bodies (MVBs), the functional consequences of enrichment in our cEVs, and EVs more broadly, remains unknown. Functional inactivation of TSG101 in mouse fibroblasts resulted in increased EGFR recycling, leading to prolonged EGF-stimulated activation of ERK1 and ERK2^55,56^. Interestingly, a recent study demonstrated that AGGF1-dependent inactivation of ERK1/2 may mediate ER stress in cardiac tissue^57^. It is therefore possible that TSG101 enrichment in cEVs could alter ERK1/2 signaling in treated normal cells, leading to activation of ER stress.

Understanding lipid heterogeneity within EV subpopulations could prove critical toward understanding internalization mechanisms and signaling events which may not be captured by more commonly used proteomic and nucleic acid characterization techniques^58,59^. We found that cEVs were enriched for lipid species esterified to a palmitic acid (16:0) moiety. Palmitic acid can induce UPR in pancreatic islet cells^40,41,60^. We have directly demonstrated that palmitic acid is capable of inducing ER stress in normal pancreatic cells at a concentration of 75 μM, to our knowledge, the lowest reported dose and relevant to the concentration quantified within cEVs. We further found that cEVs were enriched for SM lipids. The ER requires complex sphingolipids for homeostasis and altered sphingolipid metabolism has shown to be a contributor to UPR^42,43^. A sudden shock of SM species into cEV treated normal cells may alter sphingolipid homeostasis, producing the necessary conditions for ER stress and UPR activation.

Metabolomic analysis of EVs in this study has augmented the discovery of potential metabolic mediators of cancer progression^61-63^. For example, EV polar metabolite analysis identified involvement of biosynthesis pathways which interface with UPR/ER stress. tRNA charging was identified as a significantly dysregulated pathway in our metabolomics analysis/ This aligned with findings from our protein-protein interaction STRING analysis, which indicated gene expression alterations related to tRNA-aminoacylation. This is striking, as abrogated tRNA-aminoacylation has been linked to induction of ER stress and subsequent activation of the UPR^64^. Given that cEVs are enriched in multiple biomolecular classes, which directly, or potentially indirectly, regulate ER stress and protein homeostasis, and that our data show palmitic acid can partially recapitulate these findings, there could potentially be a synergistic effect across biomolecular classes. The extent to which EVs containing various micro-RNA and other RNA species have been implicated in cancer further and need to be delineated in future studies.

Our ongoing studies are investigating these potential mechanisms and how the induction of UPR/ER stress in cEV treated non-tumorigenic pancreatic cells impacts their function in the long-term. These studies may prove critical toward understanding potential molecular drivers of normal cell alteration at the earliest stages of PDAC development.

## Materials and Methods

### Cell culture

PDAC cell lines PANC-1 (CRL-1469), Capan-1 (HTB-79), SW-1990 (CRL-2172), MiaPDAC-2 (CRM-CRL-1420) were purchased from American Type Culture Collection (ATCC) and grown in modified MEM (Gibco, #A1048801) with 10% HI-FBS (Gibco, #10082147) and 1% Penicillin-Streptomycin (Gibco, #15140122). Normal pancreatic epithelial cell lines hTERT-HPNE (CRL-4023) cells were also purchased from ATCC and HPDE-H6c7 (ECA001-FP) cells were purchased from Kerafast and grown in keratinocyte serum-free media supplemented with bovine pituitary extract and recombinant epidermal growth factor (Gibco, #37010022) with 1% Penicillin-Streptomycin. The patient-derived xenograft PDAC cell lines PPCL-68 and PPCL-46 were generated as previously described^31^ and cultured in advanced MEM (Gibco, #12492013) with 5 mM L-Glutamine (Gibco, #25030081), 10% HI-FBS and 1% Penicillin-Streptomycin. All cells were grown in a 5% CO_2_ incubator at 37 °C. All cell lines were authenticated and tested by DNA fingerprinting short tandem repeat analysis (STR) at the Lombardi Comprehensive Cancer Center Tissue Culture and Shared Resource. Testing for mycoplasma and contaminants were negative for all cell lines used.

### EV isolation from conditioned media and characterization

We have submitted all relevant data of our experiments to the EV-TRACK knowledgebase (EV-TRACK ID: EV210204)^65^. cEVs were isolated from 6 PDAC cancer cell lines (PANC-1, MiaPaCa-2, Capan-1, SW-1990, PPCL-46 and PPCL-68) and 2 normal pancreatic cell lines (hTERT-HPNE and HPDE-H6c7). EV preparations were characterized using immunoblot, nanoparticle tracking analysis and cryogenic electron microscopy (methods below). We also developed a SMAD2/3/4-response element-luciferase reporter system stably expressed in a fibroblast cell line to measure the biological activity of cEV isolations (**Suppl. Fig. S8**, methods below). Once biological activity was confirmed, EVs were used for downstream experiments.

#### Cell culture conditions

When cells were ∼50 - 60% confluent, media was changed to the cells respective base media containing 10% exosome depleted FBS (Gibco, A2720801) instead of 10% HI-FBS. For normal cells, no FBS was added, and media was changed to fresh serum free media. Cells were grown for an additional 48 hours to make conditioned media. EVs were isolated from conditioned media using either differential ultracentrifugation with filtration (UC), a magnetic bead-based method, EVTrap (EVT) or size exclusion chromatography with filtration (SEC).

#### UC

Conditioned media was collected from tissue culture flasks and moved to sterile 50 mL conical tubes. Samples were spun at 1,600 x g for 20 minutes, at 4 °C. Samples were then transferred by inversion to fresh ultracentrifuge tubes (Beckman, #326823) and placed in a SW-28 (Beckman) rotor. Tubes were then spun at 10,000 x g for 20 minutes at 4 °C in an ultracentrifuge (Beckman, L8-70M). The supernatant was filtered using 0.2 μM syringe filters (Millipore Sigma, #SLGP033RB) into new ultracentrifuge tubes. Samples were placed back in a SW-28 rotor and spun at 120,000 x g for 70 minutes, at 4 °C. Supernatant was removed from tubes by inversion and the final EV pellet was re-suspended in 50 μL 1 X DPBS. Samples were stored at -80 °C until further processing.

#### SEC

Conditioned media was collected from tissue culture flasks and moved to sterile 50 mL conical tubes. Samples were spun at 2,500 x g for 10 minutes at 25 °C. Supernatants were next transferred to clean 50 mL conical tubes and again spun at 2,500 x g for 10 minutes at 25 °C. Samples were then filtered and concentrated using 100 kDa centrifugal filters (Pall Corporation, #OD100C65). Samples were spun at 3,000 x g for 30 minutes at 25 °C. Flow through was discarded and the remaining supernatant was added to filters. Samples were again spun at 3,000 x g for 30 minutes at 25 °C. This was repeated until all supernatant was filtered/concentrated. Next, concentrated media was collected and further filtered using 15 mL Amicon 100 kDa filters (Millipore Sigma, #UFC9100) by spinning at 3,000 x g for 20 minutes at 25 °C. EVs were subsequently isolated from the final concentrated media using 70 nM SEC columns (IZON, qEV2, #SP4) using an automated fraction collector, according to manufacturer’s protocol. Fractions were stored at -80 °C. Fractions containing EVs (F1-F3) were lyophilized, resuspended in 1 x DPBS, and combined to make a working stock solution of EVs.

#### EVT

Conditioned media was collected from tissue culture flasks and moved to sterile 50 mL conical tubes. Samples were spun at 1,600 x g for 20 minutes at 25 °C. Supernatant was transferred to clean 15 mL tubes. Next, media loading buffer was added to the tubes, and samples were mixed by inversion. Magnetic EVtrap beads^66^ were then added to samples and tubes were incubated for 1 hour with end-over-end rotation at 25 °C. Tubes were then placed on a magnetic separator and solution was removed. Beads were washed with media loading buffer, followed by two washes with 1 x DPBS. EVs were eluted from beads using 100 mM triethylamine (TEA) by incubation with rigorous shaking at room temperature for 10 minutes. EVs were collected, samples were lyophilized and re-suspended in 1 x DPBS. Samples were then stored at -80 °C until further use.

#### EV marker immunoblot

To investigate the expression of EV-associated proteins, we measured the protein levels of CD63, CD81, TSG101, EpCAM, ALIX, ANXA5, FLOT1, GM130 and ICAM by immunoblot using the Exo-Check Exosome Antibody Array (System Biosciences, #EXORAY210A). Protein concentration for EV samples was measured using BCA (Thermo Fisher, #23225) and 30 μg of protein was analyzed according to manufacturer’s protocol.

#### Nanoparticle Tracking Analysis (NTA)

NTA was accomplished using a NanoSight NS300 (Malvern Panalytical) equipped with a high sensitivity sCMOS camera, 531 nm laser and automatic syringe pump. 1 μL of EVs resuspended in 1x PBS was diluted to a final volume of 1 mL with 1x PBS prior to being loaded on the automatic syringe injector. Camera and detection settings can be found in **Suppl. Data 5**. Videos were captured and processed using NTA 3.3 Dev Build 3.3.104 with 3 videos of 30 seconds per measurement, per sample. Concentrations as determined by NTA were used for downstream experiments.

#### Cryogenic electron microscopy

EV samples resuspended in 1x DPBS were frozen at - 80 °C and shipped on dry ice to the Molecular Electron Microscopy Core at the University of Virginia. An aliquot of sample (∼3.5 μl) was applied to a glow-discharged, perforated carbon-coated grid (2/1-3C C-Flat; Protochips, Raleigh, NC), manually blotted with filter paper, and rapidly plunged into liquid ethane. The grids were stored in liquid nitrogen, then transferred to a Gatan 626 cryo-specimen holder (Gatan, Warrrendale, PA) and maintained at ∼180°C. Low-dose images were collected at a nominal magnification of 29,000X on a Tecnai F20 Twin transmission electron microscope (ThermoFisher Scientific, Hillsboro, OR) operating at 120 kV. The digital micrographs were recorded on a Gatan US4000 CCD or a Teitz XF416 camera.

### RNA-Sequencing

RNA was isolated from cells using the RNeasy Mini Kit (Qiagen, #74104) according to the manufacturers protocol. Library construction and sequencing was performed by Novogene Corporation (Sacramento, CA). RNA quality control (QC) was assessed using Qubit (Thermo Fisher) and Bioanalyzer (Agilent) analysis. Libraries were prepared using NEBNext Ultra II non-directional RNA Library Prep kit (New England BioLabs, # E7770S). Library quality and concentration was assessed with Labchip and qPCR. Libraries were sequenced on a Novaseq6000 (Illumina) using PE150 sequencing at a depth of 30 mm reads. Downstream analysis was performed using STAR, HTSeq, Cufflink and custom scripts. Reference genome and gene model annotation files were downloaded directly from NCBI and paired-end clean reads were aligned using STAR (v2.5). Fragments Per Kilobase Million (FPKM) counts for each gene were calculated based on the length of the gene and read counts mapped to that gene.

### Mass spectrometry solvents and standards

All solvents, including high purity formic acid (#A117-50), dichloromethane (#EW-88016-02), chloroform (#EW-80044-89), acetonitrile (#A955-4), methanol (#A456-4), isopropyl alcohol (#A461-4), and water (#W6-4) were LC-MS grade and purchased from Fisher Scientific. Debrisoquine (# D1306100MG) and taurine-d4 (#703443-100mg) were purchased from Sigma-Aldrich.

### Polar metabolite profiling

1. Sample preparation: 50 μL of EVs were transferred to glass tubes for extraction. 0.9 mL of water was added to each tube and samples were incubated on ice for 10 minutes. Next, 2 mL of methanol and 0.9 mL of dichloromethane (DCM) was added to each sample and tubes were mixed gently, but thoroughly, for 5 seconds. Samples were then incubated at room temperature for 30 minutes. Next, 1 mL of water and 2 mL of chloroform was added to each sample. Tubes were then spun at 2,000 x g for 20 minutes at room temperature. The upper aqueous layer was collected and stored at -80 °C for 5 hours. Samples were then lyophilized and re-suspended in 0.2 mL of a 1:1 mixture of acetonitrile and water containing 200 ng/mL of debrisoquine (internal standard for positive mode) and 200 ng/mL of taurine-d4 (internal standard for negative mode). Samples were centrifuged and transferred to mass spectrometry glass vials just prior to data acquisition.
2. Data acquisition: 5 μL of sample was injected onto a Kinetex 2.6 μm 100 Å 100 × 2.1 mm (Phenomenex, #00A-4723-AN) using a SIL-30 AC auto sampler (Shimazdu) connected with a high flow LC-30AD solvent delivery unit (Shimazdu) and CBM-20A communication bus module (Shimazdu) online with a QTRAP 5500 (Sciex) operating in positive and negative ion mode. Samples were resolved at a 0.2 mL/min flow rate starting with 100% of solvent A holding for 2.1 minutes and moving to 5% of solvent A over 12 minutes, holding for 1 minute before equilibrating to initial conditions over a period of 7 minutes. Auto sampler temperature was 15 °C and oven temperature was 30 °C. A binary solvent comprising of water (with 0.2% formic acid) and acetonitrile (with 0.2% formic acid) was used. Full gradient and MRM transitions that were used for quantitation are detailed in **Suppl. Data 6**. The data were normalized to internal standard area and processed using MultiQuant 3.0.3 (Sciex). Source and gas settings for the mass spectrometer were as follows: curtain gas = 35, CAD gas = Medium, Ion Spray Voltage = 2500 V in positive mode and -4500 V in negative mode, temperature = 400 °C, nebulizing gas = 60 and heater gas = 70.
3. Metabolite validations using multiple reaction monitoring-mass spectrometry (MRM-MS): this method was designed to measure palmitic acid (Sigma, #P0500), succinate (Sigma, #398055), uracil (Sigma, #U0750), ureidopropionic acid (Sigma, #94295), ornithine (Sigma, #W419001), phenyl-acetyl-l-glutamine (Sigma, #SMB00962), citrulline (Sigma, #C7629), arginine (Sigma, #A5006), and phenylalanine (Sigma, #P2126) using the QTRAP 5500 system. For calibration curves, twelve concentration points were used ranging from 1 ng/mL to 2500 ng/mL. LC-MS conditions were the same as above. For sample preparation, 25 μL of EV samples were combined with 40 μL of 35% water, 25% methanol and 40% isopropyl alcohol. Samples were plunged into dry ice for 30 seconds and heat shocked by plunging into a 37 °C water bath for 90 seconds. This was repeated for a total of 3 times. Samples were then sonicated for 30 seconds. Next, 200 μL of methanol containing 200 ng/mL of debrisoquine and 200 ng/mL of taurine-d4 was added to samples. Tubes were vortexed for 30 seconds, incubated on ice for 20 minutes and incubated at -20 °C for 20 minutes. Finally, samples were centrifuged at 13,000 x g for 20 minutes at 4 °C. Supernatant was transferred to MS vials for LC-MS analysis.

### Lipidomics profiling

1. Sample preparation: 50 μL of EVs were lysed as described above, using the heat shock method. After sonication for 30 seconds, 100 μL of ice-cold isopropyl alcohol containing lipid internal standards (full list in **Suppl. Data 6**) was added to samples. Samples were vortexed for 1 minute and incubated on ice for 30 minutes. Finally, samples were incubated at -20 °C for 2 hours to complete protein precipitation, spun at 13,000 x g for 20 minutes at 4 °C and supernatant was transferred to MS vials for LC-MS analysis.
2. Data acquisition: 5 μL of sample was injected onto an Xbridge Amide 3.5μm, 4.6 × 100 mm column (Waters Corporation, #186004868) using a SIL-30 AC auto sampler connected to a high flow LC-30AD solvent delivery unit and CBM-20A communication bus module online with a QTRAP 5500. A binary solvent comprising of acetonitrile and water (95:5) with 10 mM ammonium acetate as solvent A and 50:50 acetonitrile and water with 10 mM ammonium acetate as solvent B was used for resolution. Lipids were resolved at 0.7 mL/min flow rate with initial gradient at 100% of solvent A, shifting towards 99.9% solvent A over 4 minutes. Finally, the gradient washed with 100% of solvent B for 6 minutes and equilibrated to initial conditions of the remaining 6 minutes. Full gradient and MRM transitions can be found in **Suppl. Data 6**. Data were normalized to internal standard area and processed using MultiQuant 3.0.3. Source and gas settings were as follows: curtain gas = 30, CAD gas = Medium, Ion Spray Voltage = 5.5 kV in positive mode and -4.5 kV in negative mode, temperature = 550 °C, nebulizing gas = 50 and heater gas = 60.

### Proteomics profiling of EVs

1. Sample preparation: 50 μL of EVs were dried by lyophilization. Samples were then lysed to extract proteins using the phase-transfer surfactant (PTS) aided procedure^66^. Proteins were reduced and alkylated by incubation in 10 mM TCEP and 40 mM CAA for 10 min at 95 °C. Samples were next diluted fivefold with 50 mM triethylammonium bicarbonate and digested with Lys-C (Wako) at 1:100 (wt/wt) enzyme-to-protein ratio for 3 h at 37 °C. Trypsin was added to a final 1:50 (wt/wt) enzyme-to-protein ratio for overnight digestion at 37 °C. To remove the PTS surfactants from the samples, the samples were acidified with trifluoroacetic acid (TFA) to a final concentration of 1% TFA, and ethyl acetate solution was added at 1:1 ratio. The mixture was vortexed for 2 min and then centrifuged at 16,000 × g for 2 min to obtain aqueous and organic phases. The organic phase (top layer) was removed, and the aqueous phase was collected. This step was repeated once more. The samples were dried in a vacuum centrifuge and desalted using Top-Tip C18 tips (Glygen) according to manufacturer’s instructions. The samples were dried completely in a vacuum centrifuge and stored at -80 °C.
2. LC-MS/MS Analysis: 20% (∼1 μg) of each dried peptide sample was dissolved in 10.5 μL of 0.05% trifluoroacetic acid with 3% (vol/vol) acetonitrile containing spiked-in indexed Retention Time Standard containing 11 artificially synthetic peptides (Biognosys). The spiked-in 11-peptides standard mixture was used to account for any variation in retention times and to normalize abundance levels among samples. 10 μL of each sample was injected into an Ultimate 3000 nano UHPLC system (Thermo Fisher Scientific). Peptides were captured on a 2-cm Acclaim PepMap trap column and separated on a heated 50-cm Acclaim PepMap column (Thermo Fisher Scientific) containing C18 resin. The mobile phase buffer consisted of 0.1% formic acid in ultrapure water (buffer A) with an eluting buffer of 0.1% formic acid in 80% (vol/vol) acetonitrile (buffer B) run with a linear 60-min gradient of 6–30% buffer B at flow rate of 300 nL/min. The UHPLC was coupled online with a Q-Exactive HF-X mass spectrometer (Thermo Fisher Scientific). The mass spectrometer was operated in the data-dependent mode, in which a full-scan MS (from m/z 375 to 1,500 with the resolution of 60,000) was followed by MS/MS of the 15 most intense ions (30,000 resolution; normalized collision energy - 28%; automatic gain control target (AGC) - 2E4, maximum injection time - 200 ms; 60sec exclusion].
3. Proteomics data processing: Raw data files were searched directly against the human Swiss-Prot database updated on July 16, 2019 with no redundant entries, using Byonic (Protein Metrics) and Sequest search engines loaded into Proteome Discoverer 2.3 software (Thermo Fisher Scientific). MS1 precursor mass tolerance was set at 10 ppm, and MS2 tolerance was set at 20 ppm. Search criteria included a static carbamidomethylation of cysteines (+57.0214 Da), and variable modifications of oxidation (+15.9949 Da) on methionine residues and acetylation (+42.011 Da) at N terminus of proteins. Search was performed with full trypsin/P digestion and allowed a maximum of two missed cleavages on the peptides analyzed from the sequence database. The false-discovery rates of proteins and peptides were set at 0.01. All protein and peptide identifications were grouped, and any redundant entries were removed. Only unique peptides and unique master proteins were reported.
4. Label-free quantitation: All data were quantified using the label-free quantitation node of Precursor Ions Quantifier through the Proteome Discoverer v2.3 (Thermo Fisher Scientific). For the quantification of proteomic data, the intensities of peptides were extracted with initial precursor mass tolerance set at 10 ppm, minimum number of isotope peaks as 2, maximum ΔRT of isotope pattern multiplets – 0.2 min, PSM confidence FDR of 0.01, with hypothesis test of ANOVA, maximum RT shift of 5 min, pairwise ratio-based ratio calculation, and 100 as the maximum allowed fold change. The abundance levels of all peptides and proteins were normalized to the spiked-in internal iRT standard. For calculations of fold-change between the groups of proteins, total protein abundance values were added together, and the ratios of these sums were used to compare proteins within different samples.

### Luciferase activity assays

For measuring ER stress, a lentiviral luciferase-reporter vector for CBF/NF-Y/YY1 promoters (ER stress response elements) was purchased (Qiagen, # CLS-9032L) and HPDE-H6c7 or hTERT-HPNE cells were infected according to manufacturer’s protocol. Cells were grown for 3 weeks under selection of puromycin until a stable cell line was generated. CBF/NF-Y/YY1 activity was confirmed using tunicamycin treatment as a positive control. To determine our EV preparations were biologically active, a cancer associated fibroblast (CAF) cell line (pCAF2) expressing TGF-β responsive SMAD2/3/4 RE-Luciferase (Qiagen, #CLS-017L) was created. pCAF2 was derived from a patient-derived xenograft (PDX) mouse model of PDAC. Informed consent was obtained from all patients preoperatively under the institutional review board approved protocol IRB201600873 at the University of Florida. Following surgical resection of the patient tumor, a 2×2mm piece of tumor tissue was isolated from the specimen. This was then implanted directly into an 8-week-old female nonobese diabetic severe combined immunodeficient mouse (Jackson Laboratory, Bar Harbor, ME)^67^. Cancer associated fibroblasts (CAFs) were isolated as previously described^68^ from xenografts. Tumor tissue was fragmented into 1mm^3^ segments. Segments were placed into a 10mM CaCl2/FBS coated six-well plate. CAFs were maintained in culture comprised of Dulbecco’s Modified Eagle’s Medium-F12 (DMEM-F12), antibiotic antimycotic solution (Fisher Scientific, Waltham, MA), 10% fetal bovine serum (FBS) (Atlanta Biologicals, Atlanta, GA), and 5% CO2 at 37 °C. CAFs were passaged after they had grown to 80% confluence at 2.5 × 10^5^ per 100mm dish. Screening of many lentiviral luciferase constructs (data now shown) for downstream activated targets of cEVs revealed that SMAD2/3/4 genes were consistently significantly upregulated post cEV treatment (**Suppl. Fig S8**). Luciferase activity was confirmed using recombinant human TGFβ (R&D Systems, #240-B-010) treatment.

### Immunoblot for proteomics validation

Independent EV isolations were performed from PANC-1, PPCL-68 and HPDE-H6c7 cell lines using ultracentrifugation with filtration as described above. Protein concentration was determined using BCA and 15 μg of protein per sample was lyophilized and resuspended in sample buffer (2x LDS and distilled H2O) + 200 mM DTT (final concentration). Samples were heated at 95 °C for 5 min and run on a 4-12% Bis-Tris NuPage gel (Thermo Fisher, #NP0321). Samples were then transferred to PVDF membranes, blocked with 1% bovine serum albumin (BSA) in tris buffer saline + 0.01% tween-20 (TBST) for 1 hour. Primary antibodies were added, and membranes were incubated according to manufacturer’s protocol. Membranes were subsequently washed 3x with TBST, incubated with secondary antibodies for 1 hour at room temperature, visualized and representative images were taken. The following antibodies were purchased from ProteinTech: Anti-TMEM-59 (#24134-1-AP). Anti-TSG101 (#28283-1-AP), Anti-RALA (#13629-1-AP), Anti-UBAC2 (#25122-1-AP). Anti-RAB40C was purchased from Millipore Sigma (#07-1392).

### Palmitic acid treatment

Palmitic acid (Sigma, #P0500) was first conjugated to bovine serum albumin (BSA) for intracellular delivery. 0.1282 grams of palmitic acid was dissolved in 100% ethanol (final concentration of 500 mM) and heated at 70 °C until dissolved. 10 μL of the 500 mM palmitic acid solution was added to 990 μL of 10% BSA containing KSFM (final concentration of 5 mM). Samples were vortexed and incubated at 55 °C for 15 minutes. Vortexing and heating was repeated once more. Serial dilutions were made in 10% BSA-containing KSFM and warmed to 37 °C prior to cell treatment. 5,000 HPDE-H6c7-ERSE luciferase cells were seeded in a 24-well plate and grown for 24 hours. Cells were then treated with various concentrations of PA-BSA for an additional 24 hours before luciferase activity was analyzed.

### Caspase 3/7 activity assay

2,500 cells were seeded in a 96-well plate. After 24 hours, cells were treated with 1.00 × 10^9 cEVs and incubated for 48 hours. Caspase 3/7 activity was measured using the Caspase-Glo® 3/7 Assay (Promega, # G8090) according to manufacturer’s protocol.

### Live/dead cell immunofluorescence microscopy

5,000 cells (hTERT-HPNE or HPDE-H6c7) were seeded in a 96-well plate. After 24 hours, cells were treated with 1.00 × 10^9 cEVs and incubated for an additional 24 hours. Cells were then stained with live/dead reagents (Invitrogen, #L3224) according to manufacturer’s protocol and the number of dead cells was counted (*n* = 3 wells per condition) and quantified.

### Quantification of cell number by DAPI staining

2,500 cells were seeded in 96-well plates and allow to grow for 24 hours. Cells were then treated with 5.00 × 10^8 cEVs and grown an additional 24 hours. To determine total cell number, cells were fixed with methanol, stained with 0.1% DAPI, images were captured under a microscope and cells per treatment were calculated in ImageJ (*n* = 3 wells per condition).

### Crystal violet staining

2,500 cells were seeded in 96-well plates and allow to grow for 24 hours. Cells were treated with 5.00 × 10^8 cEVs and grown an additional 48 hours. Cells were stained with Crystal Violet (Sigma, #3886) (0.5%) and counted in ImageJ (*n* = 3 wells per condition).

### EV internalization

EVs were stained with PKH67 (Sigma, #PKH67-GL-1KT) and unbound dye was removed using Vivaspin 20, 3 kDa MWCO centrifugal filters (Sigma, #Z629456), according to manufacturer’s protocol. 10,000 hTERT-HPNE or HPDE-H6c7 cells were seeded into a 96-well plate. The next day, media was changed and 5 μg (as determined by BCA) of labeled EVs were added to each well. After 24 hours, nuclei were stained using Hoechst dye (Invitrogen, #R37605) according to manufacturer’s protocol and representative fluorescent microscopy images were captured using a BZ-X710 (Keyence) fluorescent microscope.

### Luminex cytokine assay

HPDE-H6c7 cells were seeded in a 24-well plate with 50,000 cells per well (*n* = 4 per condition). After 24 hours, media was changed, and cells were treated with 1×10^10 cEVs or nEVs. At either 2 or 4 days post initial treatment, media was collected for cytokine analysis. Media was changed every 2 days and cells were re-treated with another dose of either cEVs or nEVs. For collection, media was immediately aspirated from wells and placed into 2 mL centrifuge tubes. Media was cleared by centrifugation for 15 minutes at 1,500 x g. Supernatant was then placed into new tubes and samples were stored at -80 °C until Luminex analysis. Luminex 200 (Luminex Corp.) was used to analyze cytokines by multiplex analysis. 48 cytokines were quantified in the cell supernatant using a Bio-Plex Pro Human Cytokine Screening Panel (Bio-Rad, #12007283) according to manufacturer’s protocol. Cytokine concentrations (pg/mL) were determined by fluorescence intensity and quantification was obtained using Bio-Plex Manager software, version 6.1 (Bio-Rad).

### Data analysis and statistics

Proteomics, metabolomics, lipidomics and RNA-Seq data were pre-processed as described above. All statistical analyses were performed using custom R scripts with log transformation. Significant features were determined using FDR-adjusted (*p* < 0.05) cutoff value. Metabolomics and lipidomics data matrices were normalized using the probabilistic quotient normalization (PQN) method^69,70^. Features with quality control (QC) sample relative standard deviations (RSD) > 20% were excluded from analysis. Proteomics data was normalized to total intensity. For RNA-Seq, differential expression analysis was performed using the DESeq2 R package (1.14.1). All other reported statistical analyses were binary comparisons using Student’s two-tailed *t*-tests with homogenous variance.

## Acknowledgements

We would like to thank the Metabolomics Shared Resource, Flow Cytometry & Cell Sorting Shared Resource and Genomics & Epigenomics Shared Resource at Georgetown University, all partially supported by NIH/NCI grant P30-CA051008. We would also like to thank Dr. Kelly Dryden from the University of Virginia. Transmission electron micrographs were recorded at the University of Virginia Molecular Electron Microscopy Core facility (RRID:SCR_019031), which is supported in part by the School of Medicine and built with NIH grant G20-RR31199. Experimental schematics were created with BioRender.com. This study was supported by the American Cancer Society (IRG-92-152-17 award number AWD4470404), Georgetown Lombardi Comprehensive Cancer Center Support Grant Developmental Funds and the Ruesch Foundation to K.U and A.K.C and by the National Center For Advancing Translational Sciences of the National Institutes of Health under Award Number TL1TR001431, and a Cosmos Scholars Grant from the Cosmos Club Foundation to C.P.H. The content is solely the responsibility of the authors and does not necessarily represent the official views of the National Institutes of Health. The opinions or assertions contained herein are the private views of the authors and are not necessarily those of the Uniformed Services University of the Health Sciences, or the Department of Defense, USA.

## Author contributions

C.P.H., S.B., A.I., B.S., K.M.H., J.G.T., V.K.S. and P.P.B. were responsible for data collection. C.P.H., Y.L., and M.G. were responsible for data analysis and interpretation. C.P.H., P.P.B., and A.K.C. were responsible for drafting and editing of the manuscript as well as for the conception and design of the work.

## Competing interests

The authors declare no competing interests. The authors alone are responsible for the content and writing of this paper.

## Materials & Correspondence

Raw proteomics data has been deposited to the Proteomics Identifications Database (PRIDE) under accession number PXD028597. Raw RNA-Seq data has been deposited to the Gene Expression Omnibus (GEO) database under accession number GSE181625. All other requests for materials, data and reagents used in this study will be fulfilled by the corresponding author.

## Notes

### Competing Interest Statement

The authors have declared no competing interest.

