## Supplemental Figures for "A multi-omics approach identifies pancreatic cancer cell extracellular vesicles as mediators of the unfolded protein response in normal pancreatic epithelial cells"

**<u>Supplementary Figure S1</u>: a.** Immunoblot arrays for EV-associated protein markers. **b.** Mean diameter sizes for each EV preparation, as determined by NTA. *N* = 3 independent EV isolations per cell line. **c.** Phase contrast and fluorescent microscopy images of hTERT-HPNE cells incubated for 18-24 hours with PANC-1 or PPCL-68 cEVs. cEVs were labeled with PKH67 prior to incubation with cells.

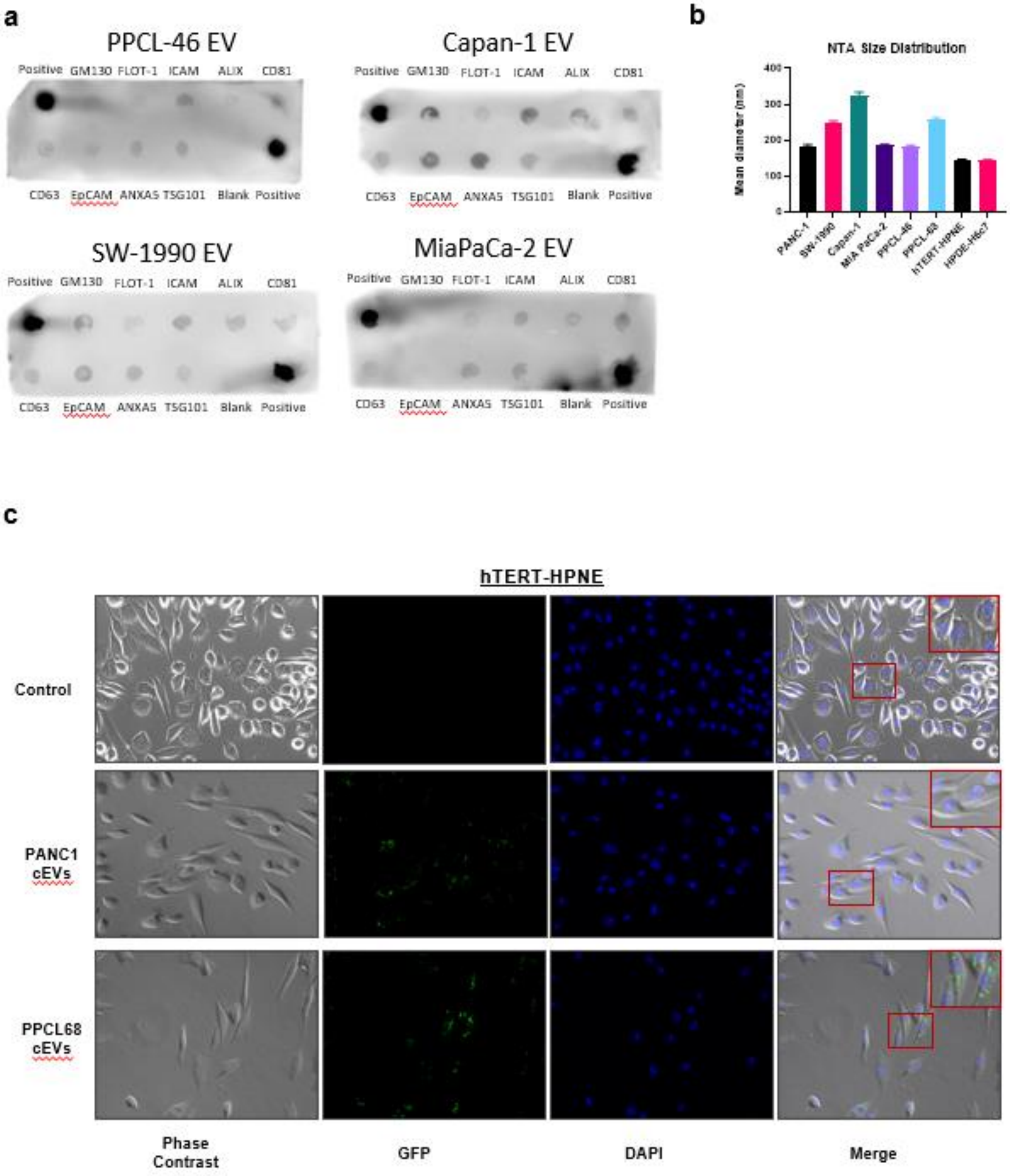
a
PPCL-46 EV
Positive GM130 FLOT-1 ICAM ALIX CD81
CD63 EpCAM ANXA5 TSG101 Blank Positive
Capan-1 EV
Positive GM130 FLOT-1 ICAM ALIX CD81
CD63 EpCAM ANXA5 TSG101 Blank Positive
SW-1990 EV
Positive GM130 FLOT-1 ICAM ALIX CD81
CD63 EpCAM ANXA5 TSG101 Blank Positive
MiaPaCa-2 EV
Positive GM130 FLOT-1 ICAM ALIX CD81
CD63 EpCAM ANXA5 TSG101 Blank Positive
b
NTA Size Distribution
Mean diameter (nm)
0
100
200
300
400
PANC-1
SW-1990
Capan-1
MIA PaCa-2
PPCL-46
PPCL-68
hTERT-HPNE
HPDE-H6c7
c
hTERT-HPNE
Control
PANC1 cEVs
PPCL68 cEVs
Phase Contrast
GFP
DAPI
Merge

**<u>Supplementary Figure S2:</u> a.** Representative cryogenic electron microscopy images from EV preparations at low (9,600x), medium (29,000x) and high (62,000x) magnifications.

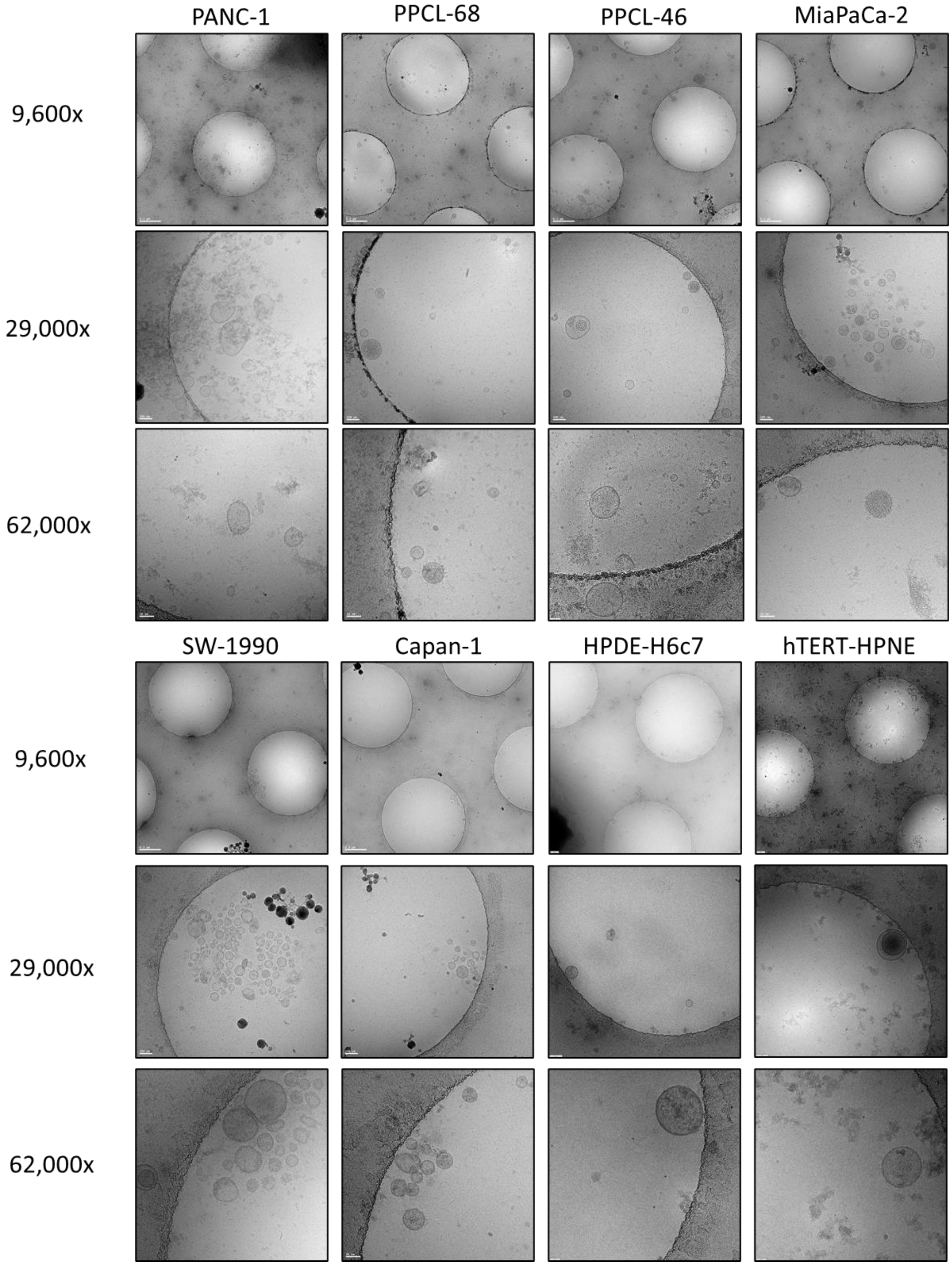

**Supplementary Figure S3: a.** STRING analysis of downregulated common DEGs between HPDE-H6c7 and hTERT-HPNE cells post cEV treatment. There was a significant increase in protein-protein interaction (PPI enrichment $p = 0.00528$) and this network was enriched for genes involved in protein refolding (adj. $p = 0.0087$). **b.** Bioplanet 2019 pathway analysis (performed through the Enrichr platform) of unique DEGs in HPDE-H6c7 cells post cEV treatment. **c.** Bioplanet 2019 pathway analysis (performed through the Enrichr platform) of unique DEGs in hTERT-HPNE cells post cEV treatment.

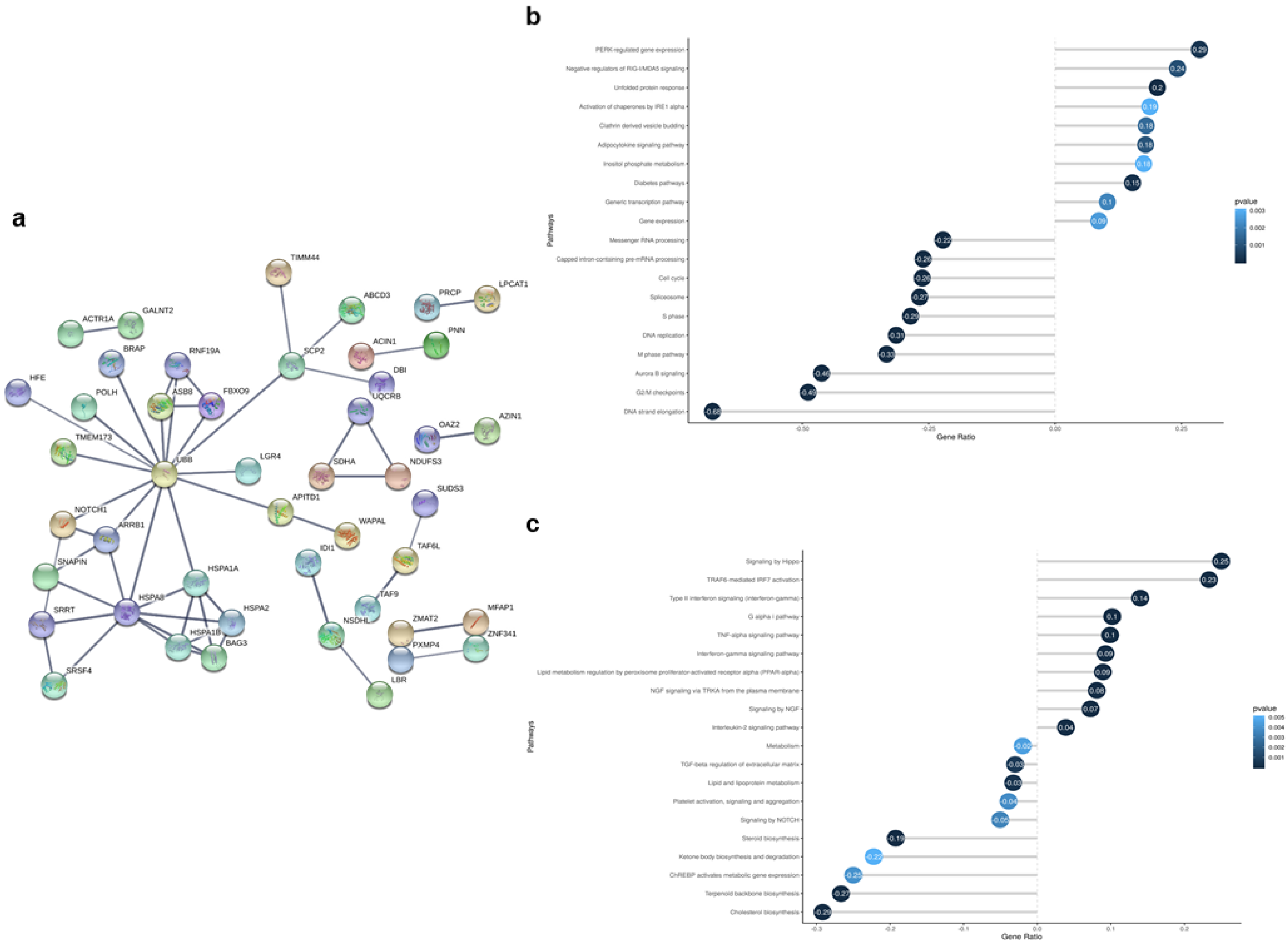
a
TIMM44
ABCD3
PRCP
LPCAT1
ACTR1A
GALNT2
BRAP
RNF19A
SCP2
ACIN1
PNN
HFE
POLH
ASB8
FBXO9
DBI
UQCRB
OAZ2
AZIN1
TMEM173
UBB
LGR4
SDHA
NDUFS3
APITD1
SUDS3
NOTCH1
ARRB1
WAPAL
TAF6L
SNAPIN
HSPA1A
IDI1
TAF9
SRRT
HSPA8
HSPA2
NSDHL
ZMAT2
MFAP1
HSPA1B
BAG3
PXMP4
ZNF341
SRSF4
LBR
b
PERK-regulated gene expression
Negative regulators of RIG-I/MDA5 signaling
Unfolded protein response
Activation of chaperones by IRE1 alpha
Clathrin derived vesicle budding
Adipocytokine signaling pathway
Inositol phosphate metabolism
Diabetes pathways
Generic transcription pathway
Gene expression
Messenger RNA processing
Capped intron-containing pre-mRNA processing
Cell cycle
Spliceosome
S phase
DNA replication
M phase pathway
Aurora B signaling
G2/M checkpoints
DNA strand elongation
Pathways
Gene Ratio
pvalue
c
Signaling by Hippo
TRAF6-mediated IRF7 activation
Type II interferon signaling (interferon-gamma)
G alpha i pathway
TNF-alpha signaling pathway
Interferon-gamma signaling pathway
Lipid metabolism regulation by peroxisome proliferator-activated receptor alpha (PPAR-alpha)
NGF signaling via TRKA from the plasma membrane
Signaling by NGF
Interleukin-2 signaling pathway
Metabolism
TGF-beta regulation of extracellular matrix
Lipid and lipoprotein metabolism
Platelet activation, signaling and aggregation
Signaling by NOTCH
Steroid biosynthesis
Ketone body biosynthesis and degradation
ChREBP activates metabolic gene expression
Terpenoid backbone biosynthesis
Cholesterol biosynthesis
Pathways
Gene Ratio
pvalue

**<u>Supplementary Figure S4:</u>** **a.** Transcription Regulatory Relationships Unraveled by Sentence-based Text Mining (TRRUST) transcriptional regulatory network analysis results, predicting transcription factors responsible for regulating genes enriched in HPDE-H6c7 cells 24 hours post cEV treatment. **b.** Transcription Regulatory Relationships Unraveled by Sentence-based Text Mining (TRRUST) transcriptional regulatory network analysis results, predicting transcription factors responsible for regulating genes enriched in hTERT-HPNE cells 24 hours post cEV treatment.

**a**

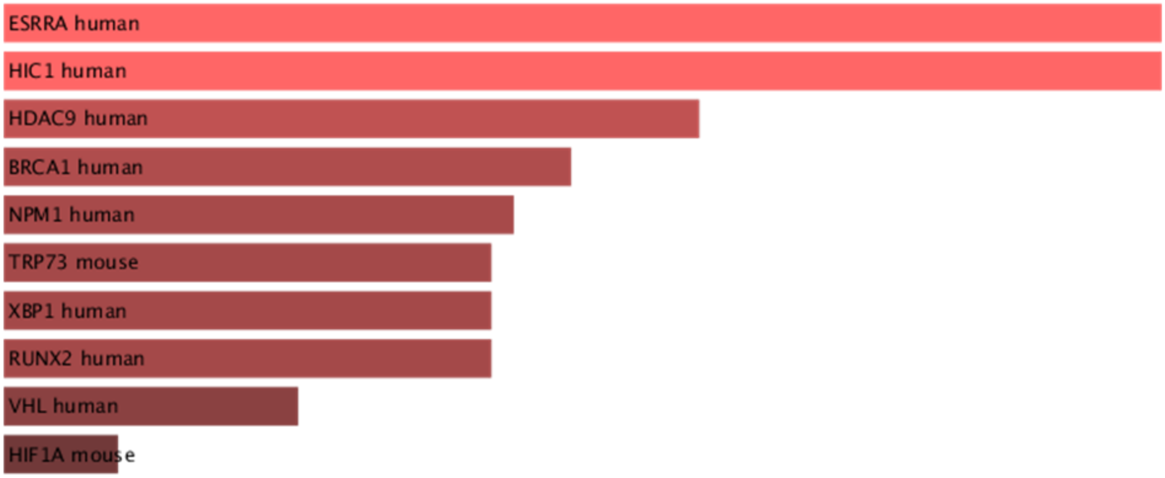


**b**

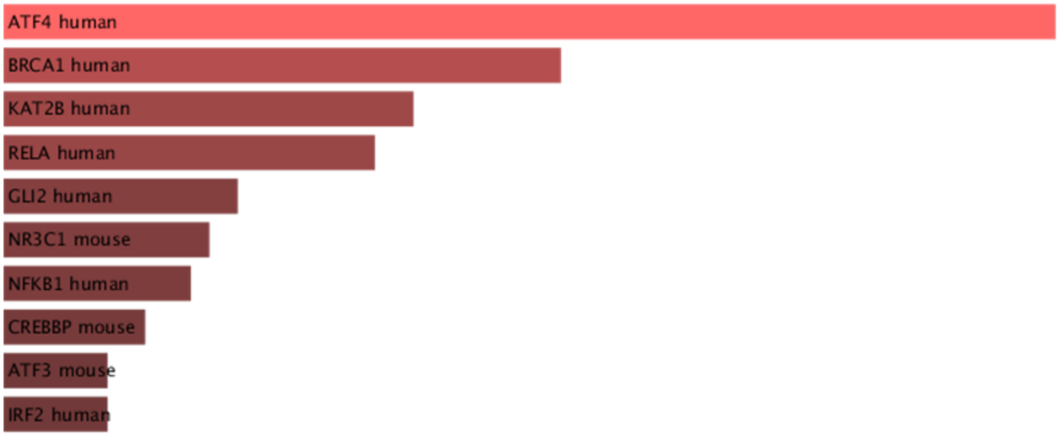

**<u>Supplementary Figure S5:</u>** **a.** Luciferase quantification showing upregulation in binding activity of the ER stress-response element (ERSE) promoter at half the dose (2.5E10 vesicles) of cEVs both 24 (top) and 48 (bottom) hours post treatment. **b.** Caspase 3/7 luciferase activity indicating no significant change in activity post cEV treatment. cEV treaments were for 24 hours while Staurosporine treatment was for 3 hours. **c.** Representative live/dead cell staining images of HPDE-H6c7 (top) and hTERT-HPNE (bottom) cells 24 hours post cEV treatment. **d.** Quantification of dead cells after live/dead staining. $N = 3$ per group. ns = $p > 0.05$, * = $p \leq 0.05$, ** = $p \leq 0.01$, *** = $p \leq 0.001$ and **** = $p \leq 0.0001$.

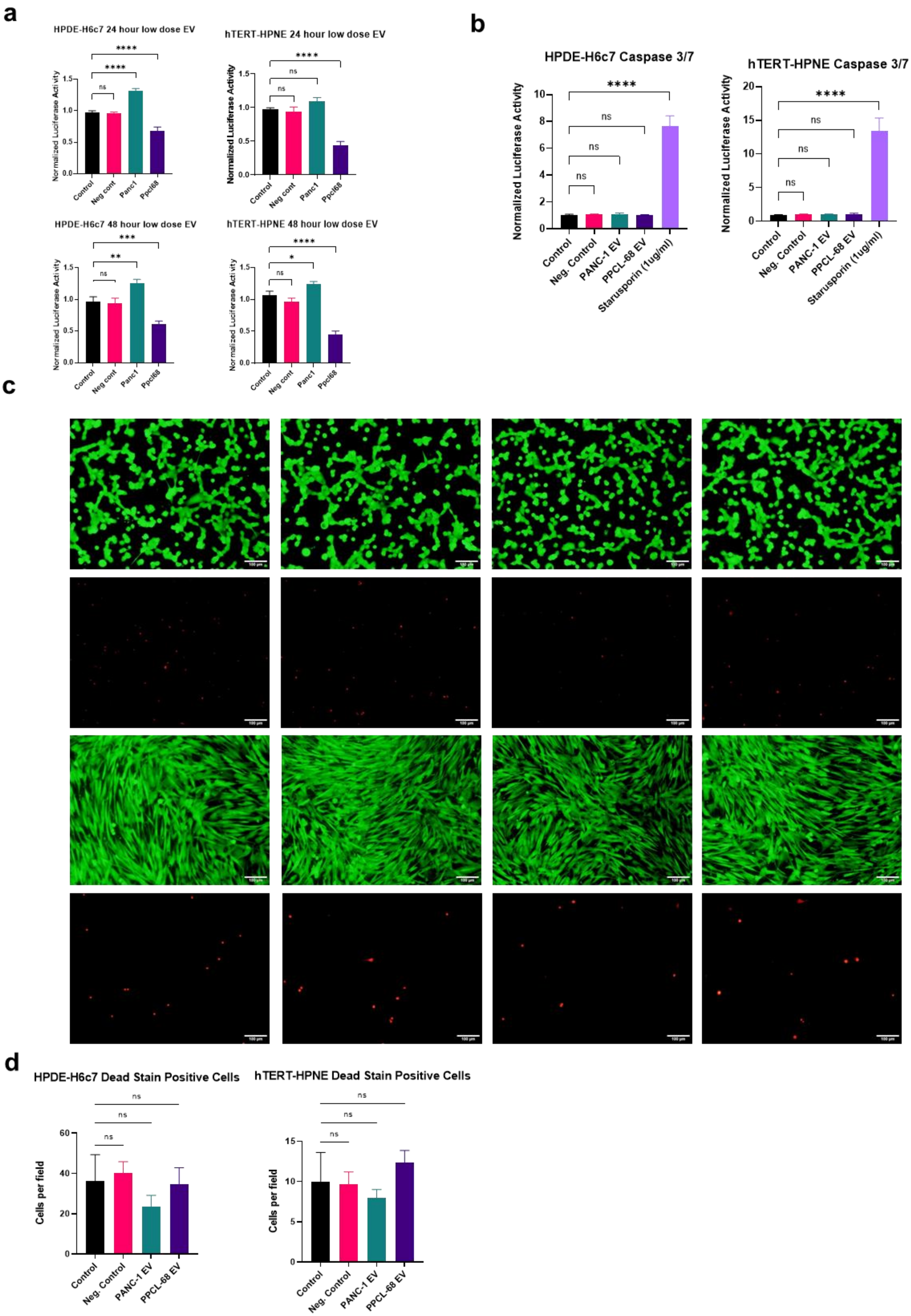

a
HPDE-H6c7 24 hour low dose EV
hTERT-HPNE 24 hour low dose EV
HPDE-H6c7 48 hour low dose EV
hTERT-HPNE 48 hour low dose EV
Normalized Luciferase Activity
Control
Neg cont
Panc1
Ppcl68
b
HPDE-H6c7 Caspase 3/7
hTERT-HPNE Caspase 3/7
Normalized Luciferase Activity
Control
Neg. Control
PANC-1 EV
PPCL-68 EV
Starusporin (1ug/ml)
c
d
HPDE-H6c7 Dead Stain Positive Cells
hTERT-HPNE Dead Stain Positive Cells
Cells per field
Control
Neg. Control
PANC-1 EV
PPCL-68 EV

**<u>Supplementary Figure S6:</u>** **a.** Quantification of nuclei counted after DAPI staining post-cEV treatment for 48 hours. $N = 3$ per group. **b.** Representative photographs from wells after staining cells with crystal violet (top) and quantifications (bottom) of cell number post-cEV treatment in hTERT-HPNE cells. ns = $p > 0.05$, * = $p \leq 0.05$, ** = $p \leq 0.01$, and *** = $p \leq 0.001$.

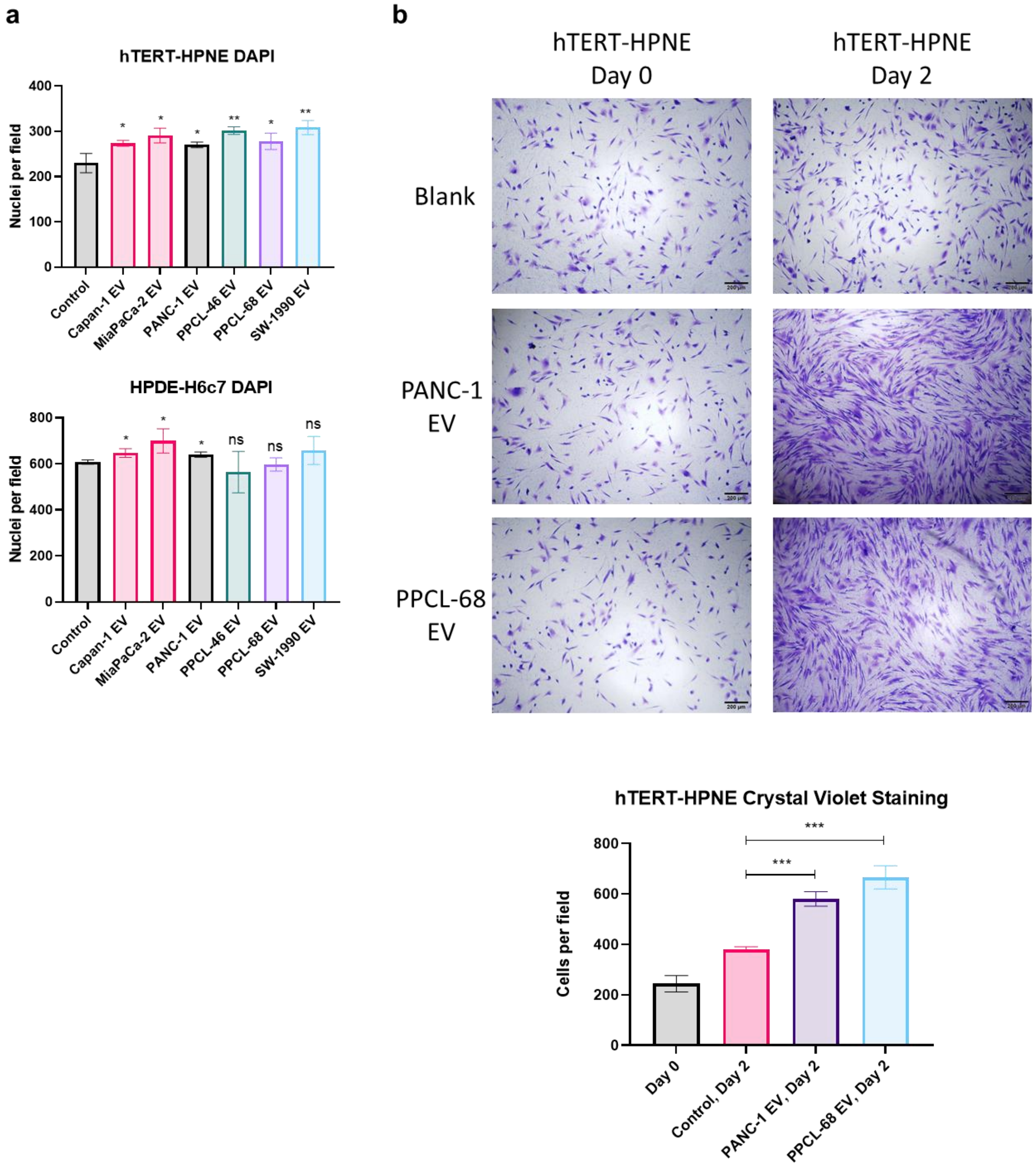

**<u>Supplementary Figure S7:</u> a.** Quantification succinate, ornithine, phenylalanine, and arginine in nEVs from HPDE-H6c7 and cEVs from PANC-1 and PPCL-68 cell lines, as quantified using MRM-MS. $N = 5$ independent EV isolations per group. ns = $p > 0.05$, * = $p \leq 0.05$, ** = $p \leq 0.01$, and **** = $p \leq 0.0001$.

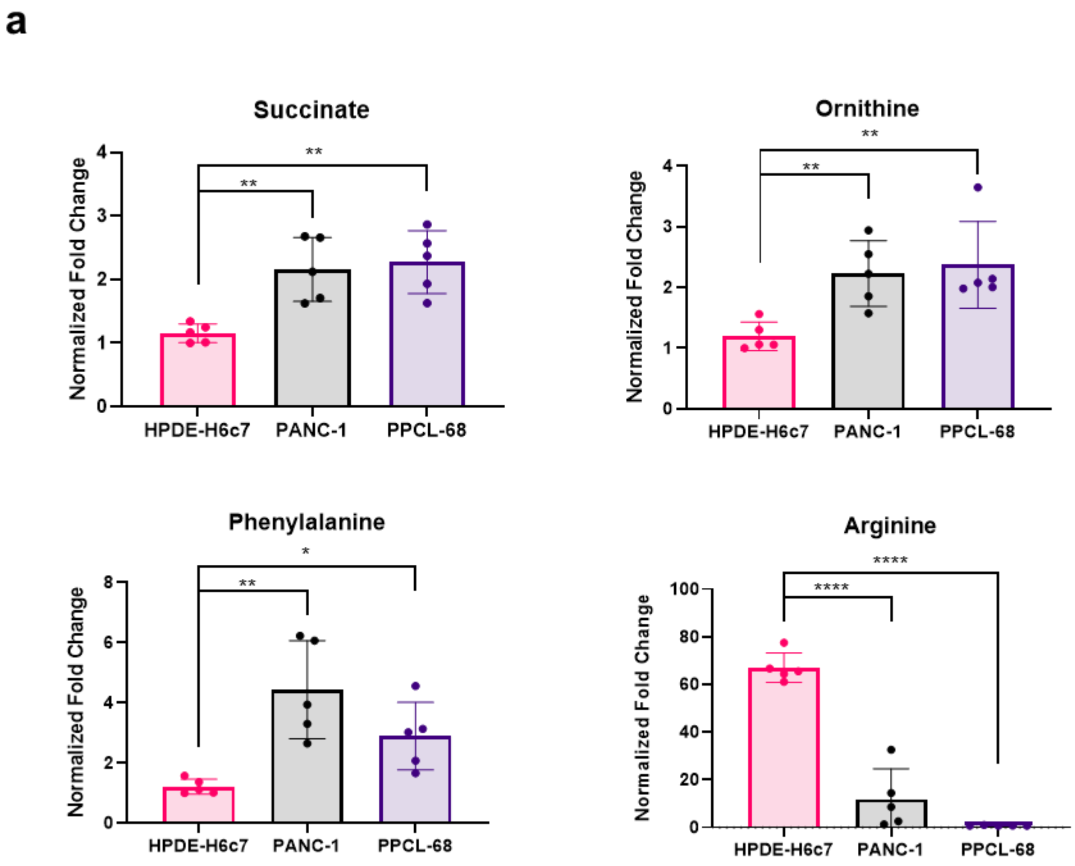

**Supplementary Figure S8:** pCAF2-SMAD2/3/4-Luciferase activity post-EV treatment, used as a measure for biological activity prior to use in downstream experiments. ns = $p > 0.05$, ** = $p \leq 0.01$, *** = $p \leq 0.001$and **** = $p \leq 0.0001$.

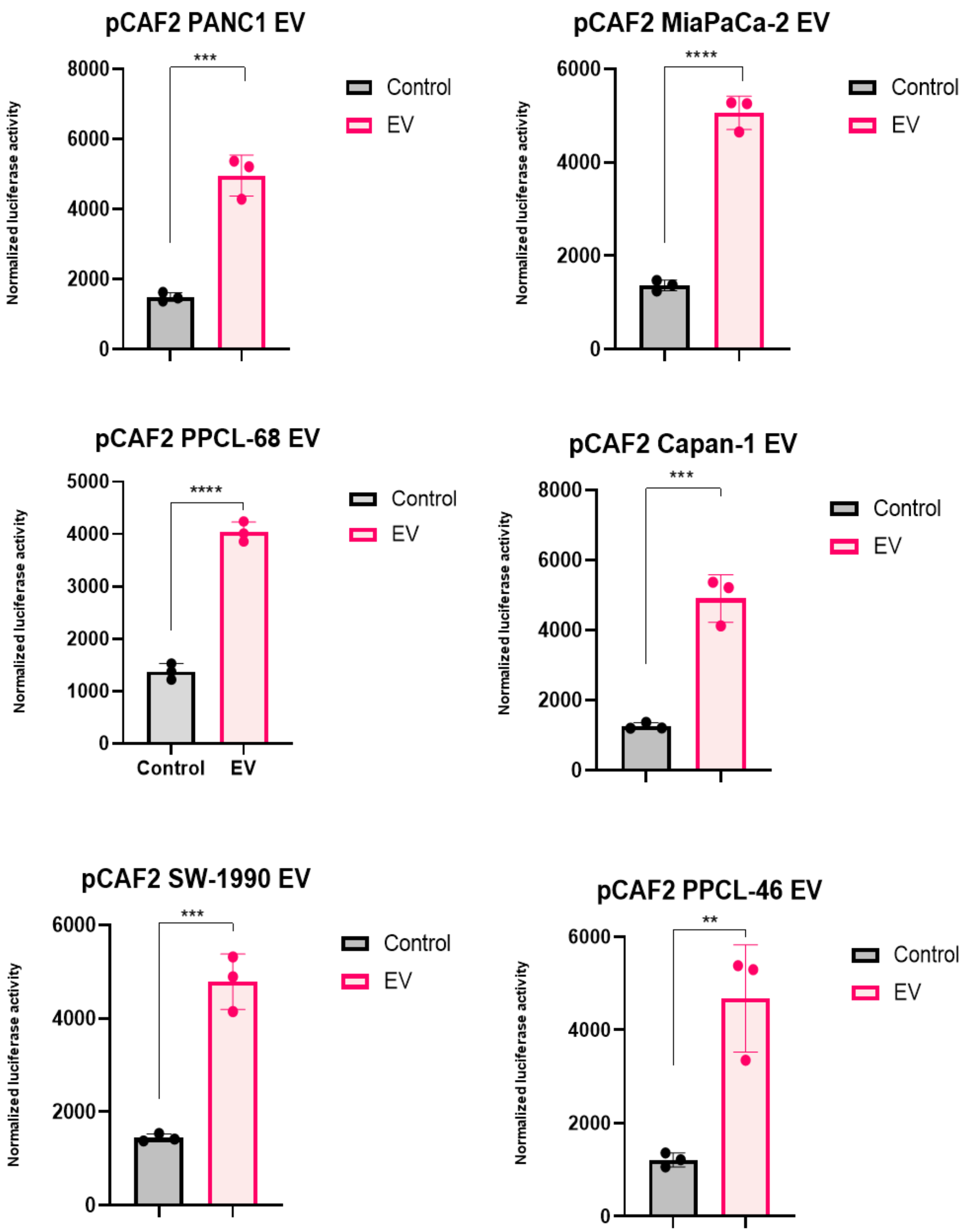
